# Reprogramming Immunosuppressive Bone Marrow–Derived Cells via CD44 Targeting Impacts Pancreatic Cancer Metastasis

**DOI:** 10.1101/2025.08.13.670111

**Authors:** Lisa-Marie Mehner, Leonel Munoz-Sagredo, Geoffroy Andrieux, David Koschut, Matthias M. Gaida, Arne Warth, Philipp Haitz, Sven Máté Treffert, Véronique Orian-Rousseau

**Affiliations:** Karlsruhe Institute of Technology, Institute of Biological and Chemical Systems – Functional Molecular Systems (IBCS-FMS), Kaiserstraße 12, 76131 Karlsruhe, Germany; Institute of Medical Bioinformatics and Systems Medicine, Medical Center - University of Freiburg, Faculty of Medicine, University of Freiburg, Freiburg, Germany; National University of Singapore, Cancer Science Institute of Singapore, Singapore 117599, Singapore; TRON Translational Oncology at the University Medical Center, JGU-Mainz, 55131 Mainz, Germany; Institute of Pathology, University Medical Center Mainz, JGU-Mainz, 55131 Mainz, Germany; Institute of Pathology, Dermatopathology, Cytology and Molecular Pathology UEGP MVZ, Wetzlar, Germany

**Author notes:** These authors contributed equally to the manuscript (Lisa-Marie Mehner, Leonel Munoz-Sagredo).

## Abstract

Pancreatic ductal adenocarcinoma (PDAC) is characterized by early dissemination and an aggressive metastatic course. In order to establish in the liver, metastatic cells require a metastatic niche providing pro-survival signals. Here, we demonstrate that bone marrow–derived cells (BMDCs) establish immunosuppressive niches in the liver that promote metastatic colonization in PDAC. Using an immunocompetent orthotopic PDAC mouse model, we show that BMDCs form clusters enriched in myeloid progenitors and display upregulation of migratory, adhesive, and immunoregulatory programs in response to tumor-derived cues. CD44 and its splice variant CD44v6 are found to be highly expressed on these cells. Hematopoietic-specific deletion of *Cd44* or *Cd44v6* using *Cd44/Cd44v6^fl/fl^;VavCreER^T2^*mice markedly impairs BMDC clustering, reshapes the BMDC transcriptome, disrupting pathways critical for migration, adhesion, and immunosuppression thereby reducing metastatic burden. Mechanistically, CD44 inhibition blocks BMDC migration toward CCL2, CCL5, and CXCL12, and impairs adhesion to VCAM-1 and fibronectin. Functionally, *Cd44*-deficient BMDCs exhibit reduced expression of immunosuppressive mediators such as Arginase 1, Ido1, and Il10, and fail to suppress T cell proliferation. Our findings position CD44 as a pleiotropic regulator of BMDC-mediated metastatic niche formation and identify it as a promising therapeutic target to disrupt the pro-metastatic microenvironment in PDAC.

## Introduction

Pancreatic ductal adenocarcinoma (PDAC) is one of the most lethal cancers^[1]^. This is mainly due to very early dissemination, producing high recurrence rates even in patients diagnosed at early stages, that underwent complete resections and received current standard-of-care combined chemotherapy^[2–4]^.

The concept of the metastatic niche and its role in metastasis initiation and maintenance has gained significant attention in recent years. In a pivotal study, Kaplan et al. demonstrated in mouse models of melanoma and lung cancer that bone marrow-derived cells (BMDCs), including hematopoietic stem and progenitor cells (HSCs/HPCs), clustered in future metastatic sites before cancer cells were detectable^[5]^. They introduced the term ‘pre-metastatic niche’ to describe the microenvironmental changes occurring around these BMDC clusters. Disrupting BMDC mobilization prevented metastasis formation, and tumor-derived factors in conditioned medium (CM) were sufficient to redirect metastatic target organs when injected into mice. These findings laid the groundwork for therapeutic strategies targeting niche formation to prevent metastasis. Over time, the niche concept evolved into a broader ‘pro-metastatic’ role, contributing to lethal systemic metastatic burden and revealing new molecular targets for disrupting niche stability.

The homing of BMDCs into tissues that have been injured appears as a biological constant. This phenomenon has been reported for the liver after experimental chronic hepatic injury with carbon tetrachloride in murine models as well as after hepatitis C virus in humans.^[6–10]^ In a similar way BMDCs have also been found in the ischemic myocardium, in the skin during wound healing, and in the lungs during usual interstitial pneumonia ^[11–13]^. Given their hematopoietic nature, BMDCs could be expected to respond to inflammatory mobilizing cues, like CXCL12 and granulocyte-macrophage colony-stimulating factor (GM-CSF) and contributing to the inflammatory process as such^[10, 14]^. However, BMDCs have also been detected in the liver in the absence of overt molecular signs of inflammation and have been shown take part in regenerative processes^[9, 15]^. Most interestingly, as for injured organs, there is a similar homing process of BMDCs towards solid tumors and to organs of future metastasis, creating the metastatic niche^[14]^. The broad multipotent and regenerative capacity of BMDCs may support metastasis development in the presence of disseminated cancer cells.

The trafficking and homing of normal HSCs/HPCs and leukemic stem cells (LSCs) in immunocompromised mice are regulated by chemokines, cytokines, proteolytic enzymes, and cell adhesion molecules (CAMs). CXCL12 promotes homing by binding CXCR4 and activating VLA-4 (α4β1 integrin), which binds VCAM1 on endothelial cells enabling transmigration, a process blocked by anti-VLA-4 antibodies. Similarly, HSC/HPCs anchorage in niches relies on CD44 binding to hyaluronan (HA), which can be disrupted by anti-CD44 antibodies. Comparable findings were observed in Acute Myeloid Leukemia (AML), where antibody treatment impaired LSC engraftment suggesting direct targeting of LSCs. We showed in an AML model that CXCL12 induces CD44/CXCR4 interaction even without HA, and that CD44 inhibition disrupts CXCL12/CXCR4 signaling, affecting stemness and survival. Thus, CD44 acts as a pleiotropic regulator in HSC/HPC trafficking.

In pancreatic cancer, we showed that targeting one molecule CD44v6 (CD44 isoforms containing the variant exon 6) on cancer cells decreased metastatic burden in several *in vivo* models, including the immunocompetent KPC (*Kras^G12D^;Trp53^R172H^;Pdx1-Cre*) model in which this intervention significantly prolonged the survival of mice already bearing advanced PDAC.^[16, 17]^ This drives attention to the role of CD44 towards the metastasis. We reasoned that targeting CD44 on BMDCs could impair metastatic niche formation in PDAC and therefore impede metastasis development.

In this paper, using RNAseq, we show that drastic changes occur in BMDCs upon growth of pancreatic cancer cells in the pancreas. Pathways related to adhesion, migration and immunosuppression get upregulated upon tumor growth. Moreover, we show that an inducible knockout of *Cd44/Cd44v6* restricted to hematopoietic cells, impairs BMDC establishment in the liver of an immunocompetent orthotopic PDAC model using KPC-derived cells. Mechanistically, we show that anti-CD44 antibodies efficiently impaired the migration of BMDCs from tumor bearing mice or cultured with tumor-factor CM towards CXCL12, CCL2, and CCL5. This treatment also blocked the adhesion of BMDCs to VCAM1 and fibronectin. Relevantly, CD44/CD44v6 loss-of-function affected the immunosuppressive transcriptomic profile and function of these BMDCs and subsequently decreased the metastatic burden. We conclude that CD44 is a plausible target to impair PDAC metastases by affecting its BMDC-driven metastatic niches.

## Results

### Hematopoietic cell clusters are detected in the liver of tumor bearing mice

We established an immunocompetent, *in vivo*, spontaneous liver metastatic model of pancreatic cancer to follow BMDC mobilization into the liver. We used a syngeneic orthotopic implantation model of the FC1245 cell line established from KPC mice implanted into the pancreas of wild-type C57BL6 mice (Figure 1A).^[16,18]^ FC1245 cells were injected into the pancreas of 6-12 weeks old mice. Two weeks after the implantation, the animals were sacrificed and the livers extracted and frozen. Cryosections of these livers exhibited clusters of hematopoietic (CD45^+^) cells in the liver parenchyma (Representative pictures in Figure 2A). The number of clusters were significantly greater in the tumor-bearing mice (TBM) compared to control mice that underwent the same implantation surgical procedure (non-tumor bearing mice or NTBM) but were injected only with PBS, in which we found few small clusters (Figure 1B). Using an anti-PDX1 staining on liver sections, we detected a few FC1245 cells in the liver at 14dpi. At 21dpi an increased number of PDX1^+^ cells within CD45^+^ cell clusters was detected (Figure 1C).

**Figure 1.**
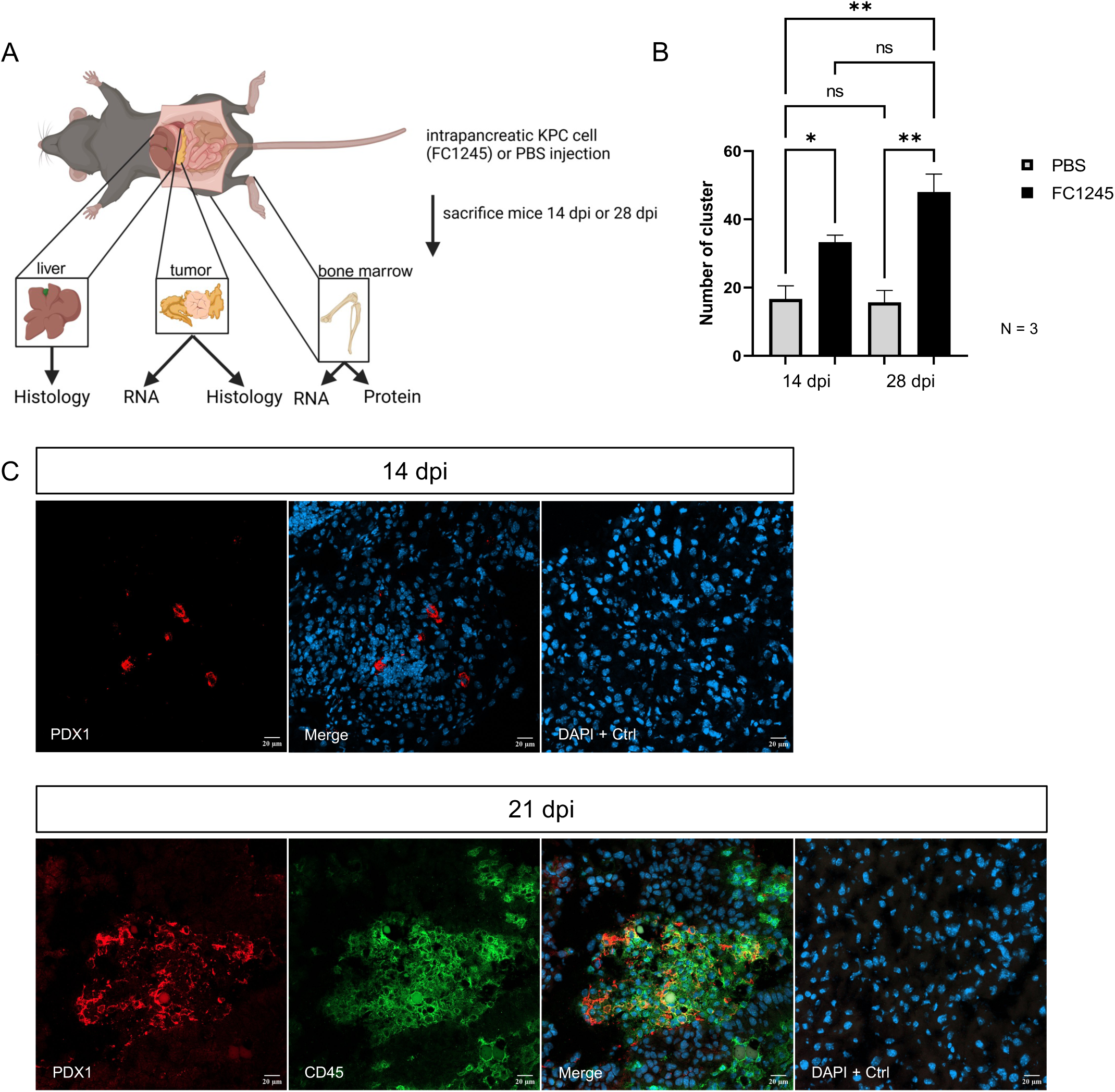
Orthotopic injection of FC1245 pancreatic cancer cells induces CD45^+^ cell clusters in the liver, which increase over time. **A** Schematic representation of the *in vivo* experimental design. FC1245 cells, derived from *LSL-Kras^G12D/+^;LSL-Trp53^R172H/+^;Pdx-1-Cre* mice, were orthotopically injected into C57BL/6 mice. PBS injections served as a control. Mice were sacrificed at 14 and 28 days post-injection (dpi). Liver tissue was analyzed histologically. Created in BioRender. Mehner, L. (2025) https://BioRender.com/poxotua **B** Immunofluorescence (IF) staining of formalin-fixed, paraffin-embedded (FFPE) liver sections from FC1245-injected and control mice. Sections were stained with an anti-CD45 antibody conjugated to FITC; isotype-matched IgG served as a control. Clusters were quantified as the mean number per animal, based on at least 3 images. **C** Immunofluorescence staining of liver sections from FC1245-injected mice at 14 and 21 dpi. At 14 dpi, sections were stained with an anti-PDX1 antibody and an Alexa546-conjugated secondary antibody. At 21 dpi, sections were stained with anti-PDX1 and anti-CD45 antibodies, followed by Alexa647- and Alexa488-conjugated secondary antibodies, respectively. Representative confocal images are shown. Scale bar: 20 µm. Statistical Analysis: Data are presented as means ± SEM. Statistical significance was assessed using one-way analysis of variance (ANOVA) with Holm-Šídák’s multiple comparisons post-hoc test. ns = not significant, *P < 0.0332, **P < 0.0021.

**Figure 2.**
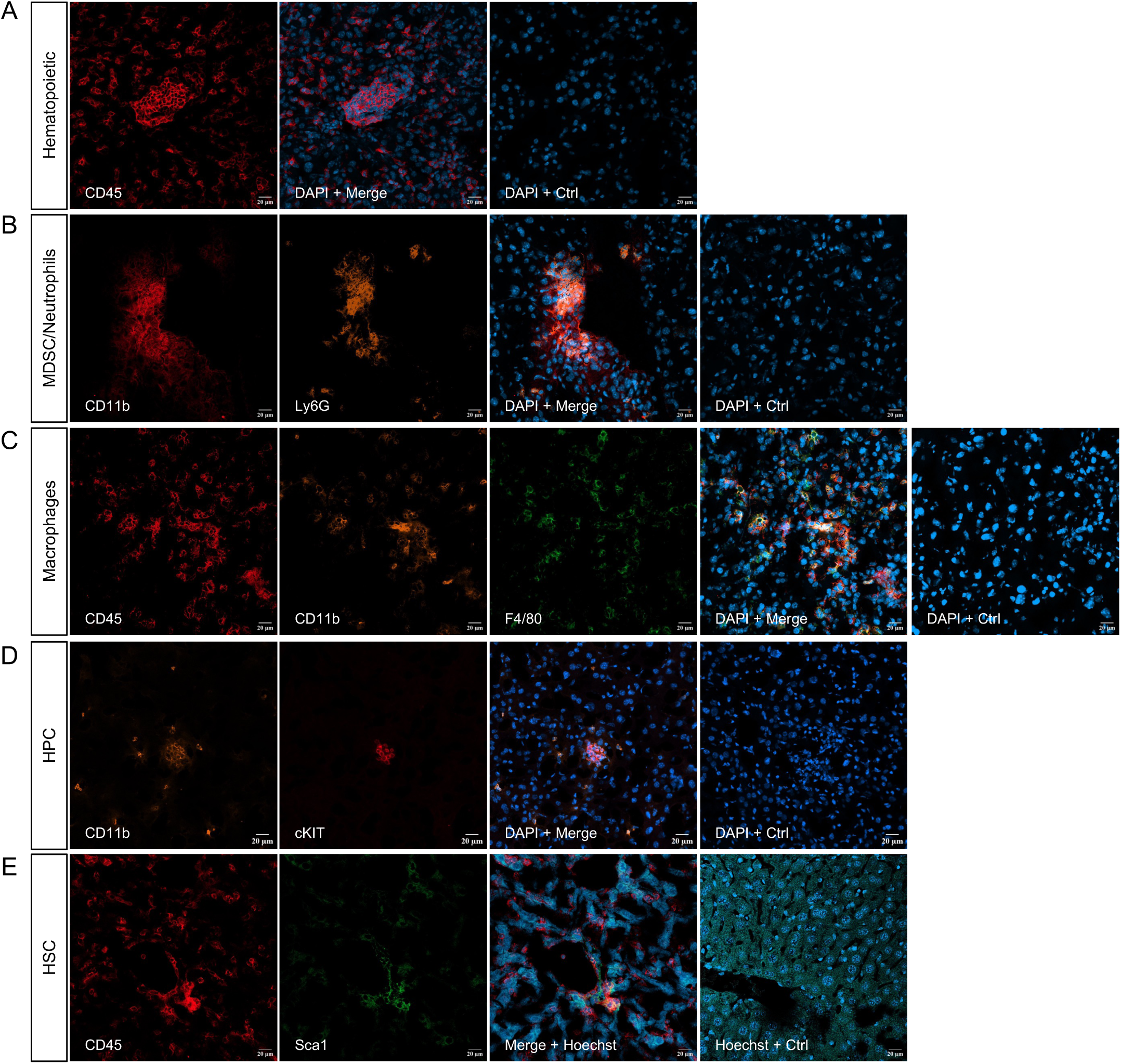
Hematopoietic cell clusters in the liver of pancreatic tumor-bearing mice contain myeloid cells and hematopoietic stem cells. Immunofluorescence of liver sections from mice orthotopically injected with FC1245 PDAC cells and analyzed at 14 dpi. **A** Staining for CD45 (APC), **B** CD11b (APC), Ly6G (PE), and DAPI **C** CD45 (APC), CD11b (PE), F4/80 (FITC), and DAPI, **D** CD11b (PE), cKIT (APC), DAPI, **E** CD45 (APC), SCA1 (FITC), and Hoechst. Isotype-matched controls were included. Confocal microscopy using the Zeiss Confocal Microscope LSM 800.; scale bar: 20 μm.

Further characterization of the CD45^+^ clusters (Figure 2A) showed that they were formed mainly by myeloid cells (CD11b^+^) (Figure 2B). Some of these cells expressed the neutrophil and granulocytic-myeloid-derived suppressor cells (G-MDSC) marker Ly6G. We additionally detected cells expressing the macrophage marker F4/80 (Figure 2C). Since the liver parenchyma is populated with Kupffer cells that also express CD11b and F4/80 and to distinguish these liver resident macrophages from incoming hematopoietic cells, we stained the same cryosections of livers from TBMs with antibodies against markers of undifferentiated hematopoietic cells present in the BM in normal conditions. A proportion of the clusters were indeed formed by c-kit^+^/Sca-1^+^ (stem cell antigen-1) cells, absent in the Kupffer cells scattered around them (Figure 2D and E). c-kit (CD117) is expressed in HSC and myeloid progenitors, while Sca-1 is confined to the HSC compartment, suggesting that the liver clusters in this model were formed by BMDCs.^[19]^ We found no components of the lymphatic lineages forming part of the hematopoietic cell clusters in the livers of TMB.

### BMDC respond to developing pancreatic tumor

The formation of pre-metastatic niches involves the mobilization and homing of BMDCs to other organs. BMDCs respond to signals coming from the tumor^[5, 20]^. We measured signals such as cytokines and growth factors released by the tumor as well as the expression of various membrane receptors on BMDCs on the other hand. Compared to normal pancreatic tissue, pancreatic tumors in our model showed high mRNA levels of secreted factors known to play an important role in PDAC, as TGF-β, PDGFα, VEGF or HGF^[21–23]^. Given the drastic increase in *Tgfb* isoforms, it suggests a major immunosuppressive role of TGF-β in our mouse model. Relevant to cell mobilization we also found increased expression of GM-CSF as well as chemokines like CCL2, CCL5, and CXCL12. As expected, mRNA levels of proinflammatory cytokines, interleukins 1β and 6 (IL1β, IL6), tumor necrosis factor α (TNFα) were also significantly increased in tumor tissue. We did not find an increased expression of epidermal growth factor (*Egf*), granulocyte colony-stimulating factor (*Gcsf*) or the chemokine *Cxcl7* (Figure 3A). Of note, interferon gamma (*Infγ*) transcription was only slightly upregulated amidst an elevation of the other pro-inflammatory cytokines mentioned. Simultaneously, we observed changes in the gene expression profile of BMDCs from TBM compared to NTBM. Supplementary Figure 3A presents heatmaps from the Gene Ontology analysis, showing the top 50 most significantly (A) upregulated biological processes in BMDCs from PDAC TBM versus NTBM. Statistical significance was determined using adjusted p-values. The greater number of significantly upregulated processes compared to downregulated pathways is displayed in Supplementary Figures 3A and 3B. All 50 upregulated processes meet the threshold of an adjusted p < 0.05, while only 6 of the downregulated processes reached statistical significance. Among the upregulated processes, 31 are immune-related (▴). In contrast, only 2 of the 6 significantly downregulated processes were linked to immune function. These findings indicate that BMDCs in PDAC-bearing animals undergo substantial functional changes, predominantly involving immune-related pathways, likely driven by tumor-derived factors. Concentrating on leukocyte-dependent pathways, we categorized these genes into three groups: leukocyte-associated adhesion (●), leukocyte-associated migration (♢) and leukocyte-mediated immunity (▴) (Supplementary Figure 3C). This suggests, that in response to the tumor, BMDCs alter biological processes involved in metastatic niche formation.

**Figure 3.**
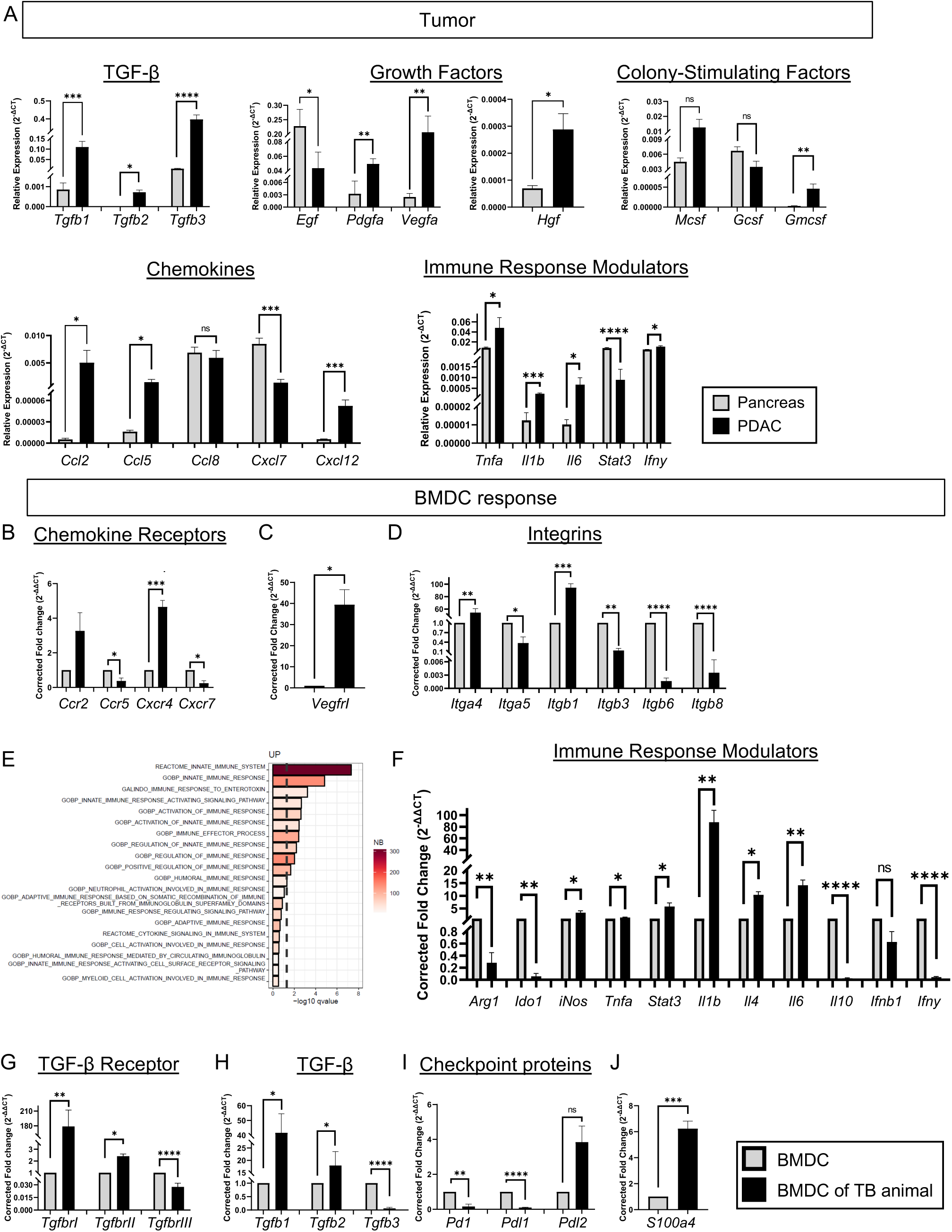
Pancreatic tumor displayed increased expression of growth factors, chemokines and inflammatory mediators, while bone marrow-derived cells (BMDC) increased the expression of the respective receptors. FC1245 cells were injected orthotopically into C57BL/6 mice. At 14 dpi, primary tumor was extracted and BMDCs were isolated from the tibia and femur. **A** Expression pattern of specific genes from the tumor was calculated as relative expression (2^−ΔCT^) and compared to healthy pancreatic tissue. *Gapdh* and *bactin* were used as reference genes for normalization. **B-D, F-J** RT-qPCR analysis of genes related to chemokine receptors and integrins expressed by the tumor compared to healthy pancreas. The corrected fold change (2^−ΔΔCT^) was calculated relative to bone marrow derived cells of non-tumor bearing C57BL/6 mice. *Gapdh* and *bactin* were used as reference genes. **E** RNA of BMDCs was subjected to bulk RNA-seq (n=5 per group). Top 20 upregulated immunity-associated GO terms, ranked by –log₁₀ (adjusted P-value) are represented; the color scale reflects the number of genes in the GSEA leading edge. Data are means ± SEM. Statistical significance was determined using the one-sample t-test; ns=not significant, *p-value<0.0332; **p-value<0.0021; ***p-value<0.00021; ****p-value<0,0001. Abbreviations: PDAC = pancreatic ductal adenocarcinoma; BMDC = bone marrow-derived cells

As migration of BMDCs is primarily mediated by chemokines, we analyzed the expression of chemokine receptors on BMDCs by qPCR (Figure 3B)^[24]^. The corrected fold change of BMDCs of TBM was normalized to BMDCs isolated from healthy animals of the same strain. Expression of the chemokine receptors *Ccr2* and *Cxcr4* was increased in response to PDAC, while *Ccr5* and *Cxcr7* expression was downregulated. Therefore, the enhanced expression of *Ccl2* and *Cxcl12* in the tumor (Figure 3A) correlated with an increased expression of the respective receptors *Ccr2* and *Cxcr4* in BMDC. Interestingly, CCL2 preferably interacts with CCR2 and CXCL12 exclusively binds to CXCR4, whereas most chemokines and receptors have several interaction partners. Given that CCL5 interacts with CCR1, CCR3 and CCR5 and CCR5 interacts with CCL3, CCL4, CCL5 and CCL8, elevated levels of CCL5 in the primary tumor and decreased expression of CCR5 in BMDCs are not contradictory. We have further detected an elevated expression of *VegfrI* (Figure 3C), which was demonstrated to be an essential receptor for BMDC infiltration^[5]^. Integrins α4 and β1 (VLA-4), relevant to BMDC adhesion at distant sites, were upregulated at the mRNA level in BMDCs from TBM, while other integrins (*Itga5, Itgb3, Itgb6, Itgb8*) were downregulated (Figure 3D). Given that most of the top 50 upregulated pathways were immune-related (Supplementary Figure 3), the top 20 immune-associated processes were further analyzed (Figure 3E). These revealed concurrent upregulation of both innate immune response pathways and regulatory processes (e.g., regulation of immune/innate immune response). qPCR confirmed a drastic increased expression of inflammatory mediators *Il1b*, *Il4*, *Il6*, a moderate upregulation of *iNos*, *Tnfa* and *Stat3* and decreased expression of immunosuppressive mediators *Arg1*, *indoleamine 2,3 -dioxygenase* (*Ido1*), *Il10*, *Ifnb1* and *Ifny* in tumor-exposed BMDCs (Figure 3F). Given the central immunosuppressive role of TGF-β in PDAC, expression of its receptors (Figure 3G) and isoforms (Figure 3H) was assessed^[25]^. BMDCs from TBM showed increased *TgfbrI/II* and *Tgfb1/2*, with reduced *TgfbrIII* and *Tgfb3*, consistent with autocrine TGF-β signaling. Checkpoint markers *Pd1* and *Pdl1* were significantly reduced, while *Pdl2* showed a non-significant trend toward upregulation (Figure 3I). Lastly, *S100a4*, a marker of pathological inflammation, was strongly upregulated in response to tumor-derived signals (Figure 3J).

### *Cd44* knockout in BMDCs impairs hematopoietic cell cluster formation in the liver and PDAC metastases

We have previously shown that CD44v6 signaling on cancer cells plays a major role in pancreatic cancer development.^[16]^ Moreover, CD44 isoforms act as co-receptors for multiple receptors upregulated in BMDCs following pancreatic cancer cell injection (Figure 3)^[26]^. Therefore, we investigated whether CD44 expression could be detected in BMDC clusters. Immunofluorescence analysis revealed strong CD44 expression within these clusters (Figure 4A) together with CD45. CD44v6 was also found expressed in the clusters (Figure 4B). Moreover, mRNA levels of CD44/CD44v6 were significantly higher in BM cells extracted from TBM compared to BMDCs of NTBM (Figure 4C). Quantification of CD44 variant exons relative to the CD44 standard isoform (CD44s) further confirmed an increased expression of most variants in BMDCs from TBM (Supplementary Figure 4).

**Figure 4.**
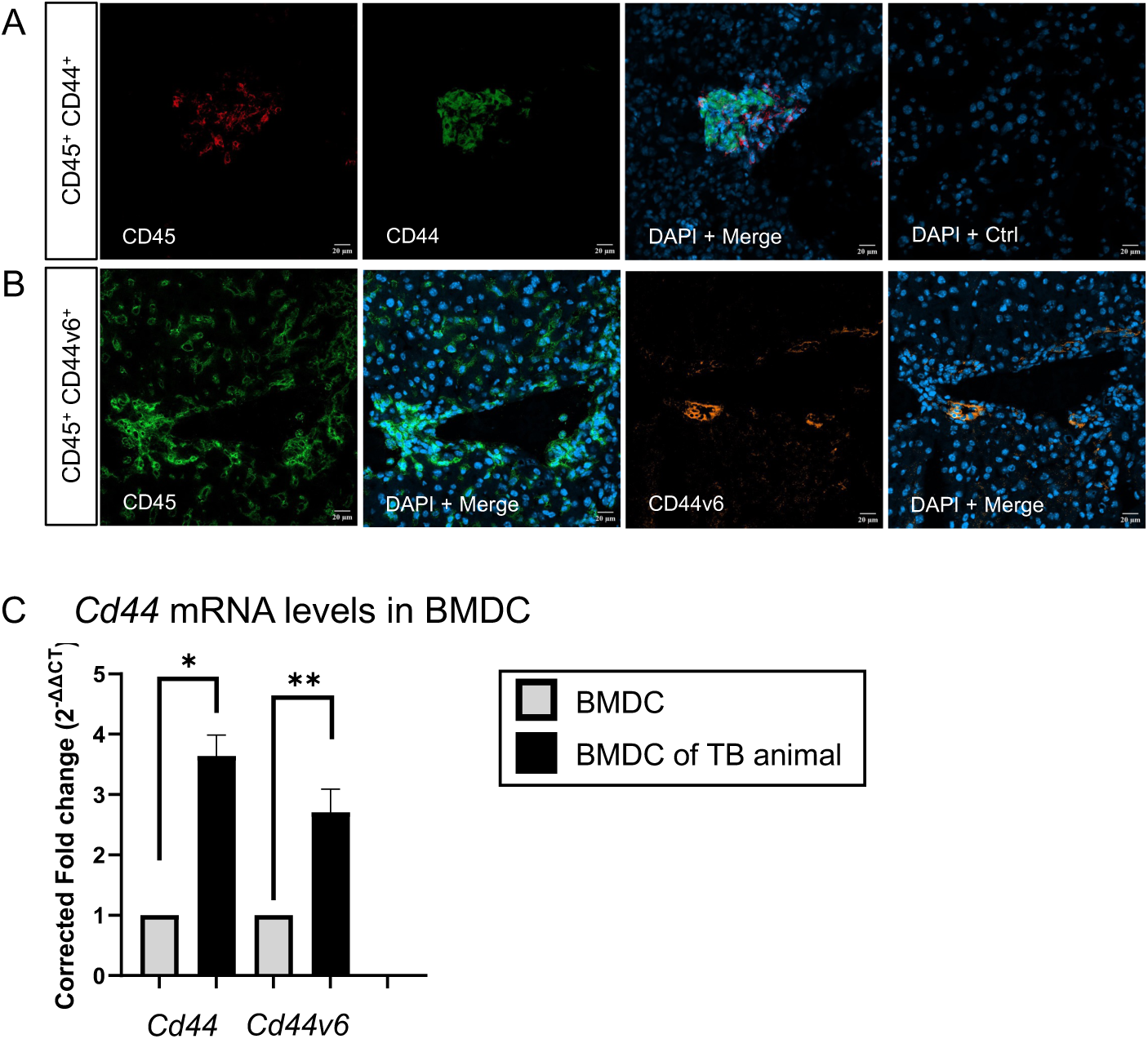
CD44 and CD44v6 are expressed on hepatic hematopoietic clusters and upregulated in bone marrow-derived cells (BMDC) from tumor-bearing mice. FC1245 pancreatic cancer cells were orthotopically injected into mice. At 14 dpi, livers were processed for histology and BMDCs were isolated from tibia and femur. **A** Immunofluorescence of liver sections stained for CD45 (APC) and CD44 (FITC); nuclei were counterstained with DAPI. Isotype controls were included. **B** Serial liver sections stained for CD45 (primary only) or CD44v6 (primary only), followed by secondary antibodies (Alexa488 for CD45, Alexa546 for CD44v6); CD45 staining serves as a specificity control for v6. Nuclei were counterstained with DAPI. **C** RT–qPCR analysis of *Cd44* (all isoforms) and *Cd44v6* in BMDCs; expression calculated as fold change (2^−ΔΔCT^) relative to BMDCs from non-tumor-bearing mice, normalized to *Gapdh*.

To explore the functional role of CD44 in BMDC cluster formation and metastasis, we utilized *Cd44^fl/fl^/Cd44v6^fl/f^*^l^ mouse models (C57BL6 background) crossed with a transgenic mouse line expressing an inducible Cre-ER^T2^ under the vav promoter, which is exclusively active in hematopoietic cells across all lineages^[27, 28]^. EYFP was used as a reporter, as Cre activity excises a lox-STOP-lox sequence in front of the EYFP locus. The knockout of *Cd44* was confirmed *in vitro* by administration of 4-hydroxytamoxifen and subsequent flow cytometry as well as assessment of the corrected fold change of *Eyfp* expression (Supplementary Figure 5A and B). The knockout of *Cd44v6* was confirmed by qPCR analysis for *in vivo* and *in vitro* experiment (Supplementary Figure 5C). In both animal models, primary tumor volume showed a trend towards a decreased volume (Supplementary Figure D).

In mice with *Cd44* deletion in BMDCs also designated *Cd44^ΔHem^*mice, we observed a marked reduction in the number of CD45^+^ clusters (Figure 5A left panel). Similarly, when *Cd44v6* was specifically deleted via tamoxifen administration in *Cd44v6^fl/fl^;VavCre-ERT2* (*Cd44v6^ΔHem)^* mice, the number of CD45^+^ clusters in the liver was significantly reduced (Figure 5A right panel). To assess the impact of CD44/CD44v6 deletion on metastasis, we quantified metastatic lesions in liver sections from these mice. Notably, the loss of *Cd44/Cd44v6* in bone marrow cells led to a significant decrease in the amount of metastases (Figure 5B). These findings strongly suggest that CD44/CD44v6 play roles that are important for the early formation of BMDC clusters during the metastatic progression of FC1245 cells.

**Figure 5.**
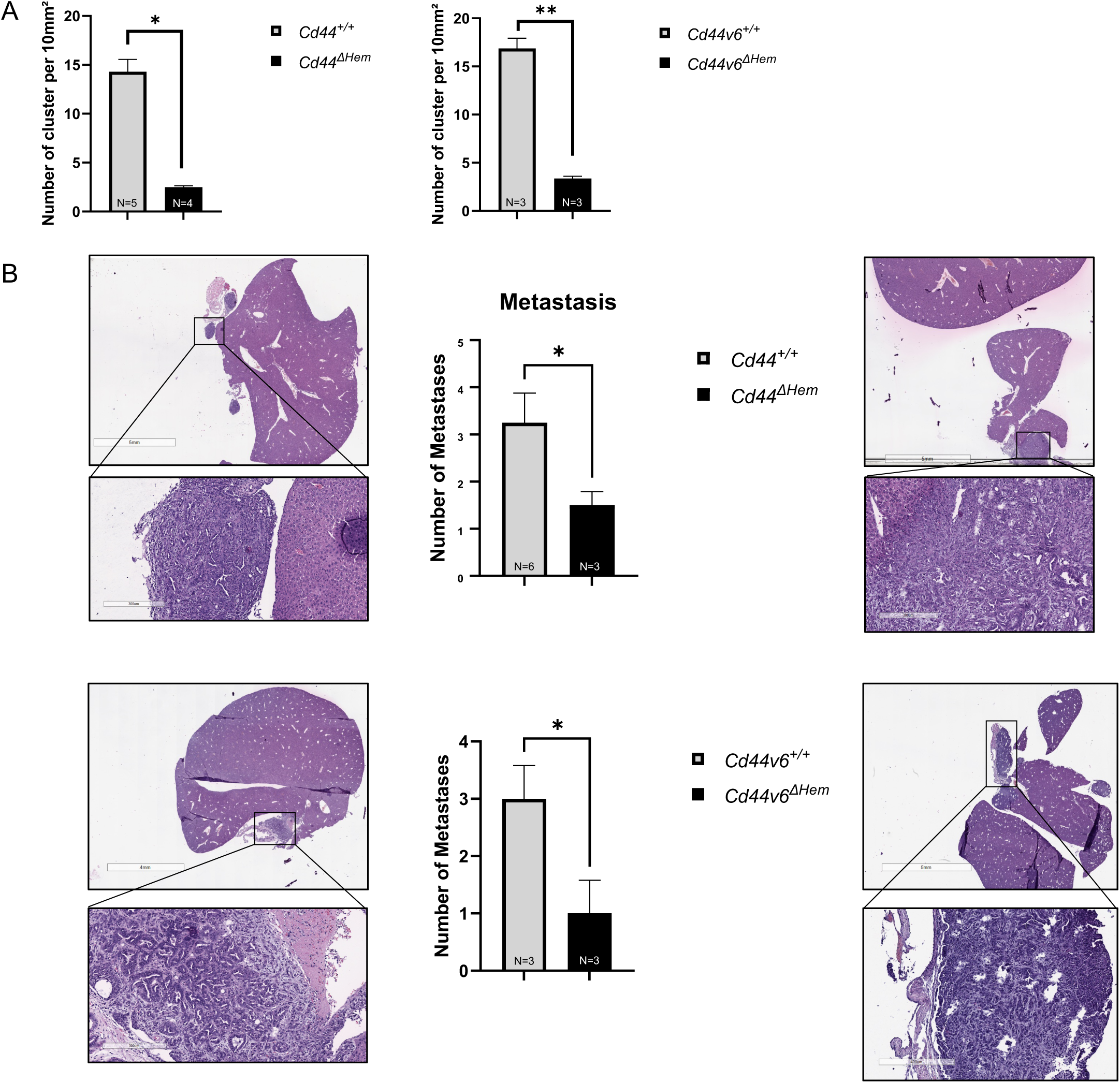
Hematopoietic-specific deletion of *Cd44* or *Cd44v6* reduces hepatic immune cell clusters and metastatic burden in pancreatic cancer. *Cd44*^fl/fl^;*VavCreER^T2^*,*RosaEYFP* (*Cd44^ΔHem^*) and *Cd44v6^fl/fl^;VavCreER^T2^;RosaYFP* (*Cd44v6^ΔHem^*) mice and their respective littermate controls (*Cd44*^+/+^, *Cd44v6*^+/+^) received tamoxifen intraperitoneally for 5 consecutive days. FC1245 pancreatic cancer cells were orthotopically injected one week later (day 0). **A** 14 dpi. Quantification of CD45^+^ clusters per mm² across serial liver sections at multiple depths (≥10 sections per animal); area measured using FIJI. **B** 21 – 28 dpi. H&E staining of FFPE liver sections. Metastases were identified by digital microscopy. Representative images are shown. Scale bar: 5 mm.

### RNAseq analysis reveals a role of CD44 expressed on BMDCs in the regulation of the immune response

To investigate the impact of CD44 deletion on BMDC function, bulk RNAseq was performed on 5 biological replicates of *Cd44^ΔHem^* mice. BMDCs from *Cd44^ΔHem^* TBMs were compared to those from *Cd44^+/+^*controls (Figure 6). GO analysis of the top 50 altered biological processes showed significant upregulation of 14 processes (Figure 6A), while all 50 downregulated processes reached statistical significance (Figure 6B). Notably, most downregulated pathways were associated with metastatic niche formation, including one related to migration (♢), six to adhesion (●), and 25 to immune responses (▴). These drastic changes were mediated by the upregulation of 12 genes and downregulation of 17 genes (Figure 6C).

**Figure 6.**
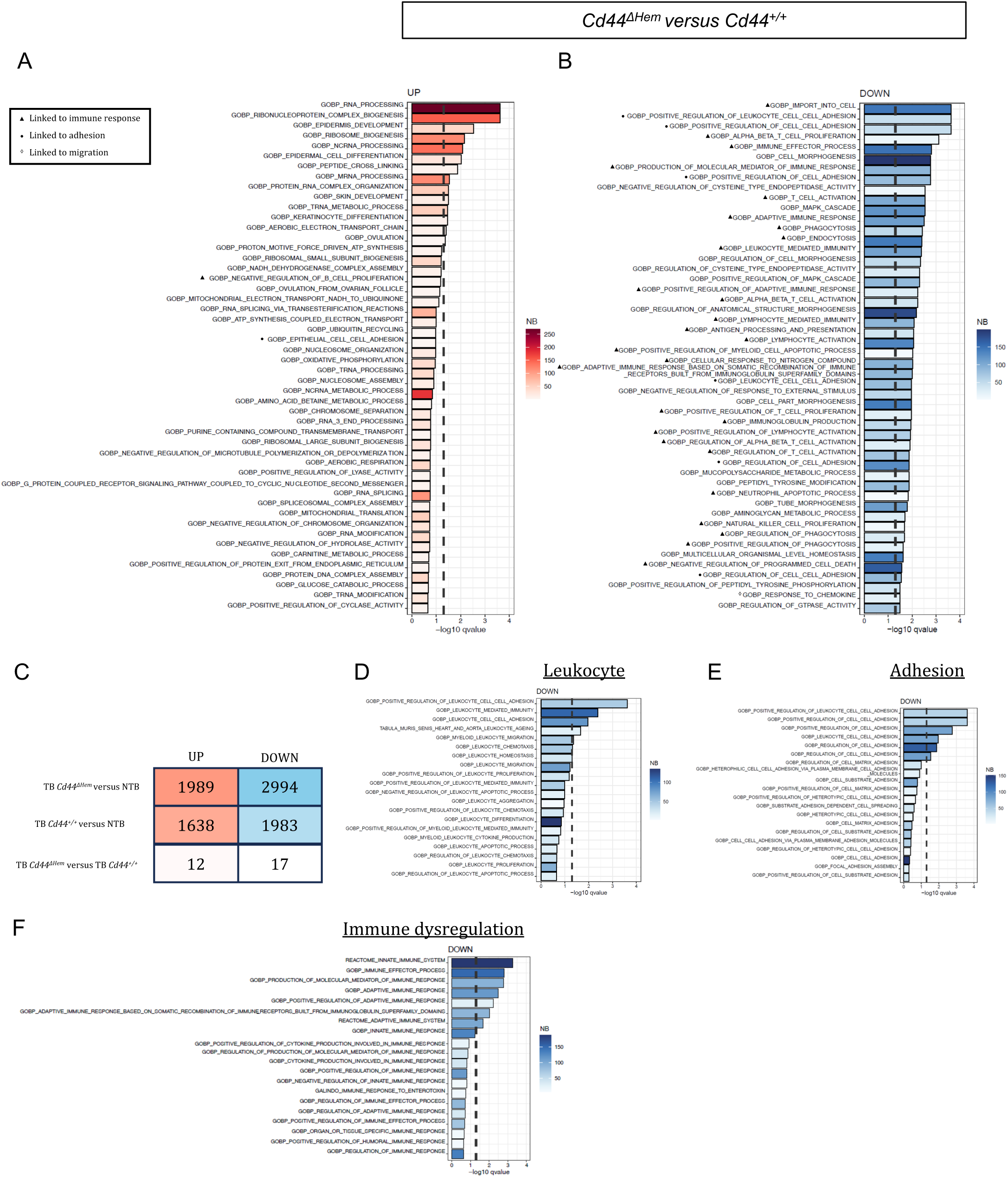
Hematopoietic-specific deletion of *Cd44* suppresses migration-, adhesion- and immune-related gene expression programs in tumor-bearing mice. FC1245 PDAC cells were orthotopically injected into *Cd44^+/+^*and *Cd44^ΔHem^* mice. At 14 dpi, BMDCs were isolated from tibia and femur and analyzed by bulk RNA sequencing (*n* = 5 per group). **A,B** Top 50 up- and downregulated Gene Ontology biological processes in *Cd44^ΔHem^* versus *Cd44^+/+^* tumor-bearing mice. Terms are ranked by –log₁₀(adjusted *P*-value); the color scale reflects the number of genes in the GSEA leading edge. **C** Heatmap showing differentially expressed genes (adjusted *P* < 0.05) across conditions: non-tumor-bearing, tumor-bearing *Cd44^+/+^* and tumor-bearing *Cd44^ΔHem^* mice. **D,E,F** Bar plots showing the top 20 downregulated terms related to: **D** leukocyte migration, **E** adhesion, and **F** immunity in *Cd44^ΔHem^* versus *Cd44^+/+^* tumor-bearing mice. Terms are sorted by –log₁₀(adjusted *P*-value); color scale represents the number of genes in the leading edge of the Gene Set Enrichment Analysis (GSEA)

To further investigate the impact of *Cd44* deletion on biological pathways, the top 20 downregulated processes related to leukocyte function (Figure 6D), adhesion (Figure 6E), and immunity (Figure 6F) were analyzed. Among the leukocyte-associated processes, 2 biological processes were related to adhesion, 5 to migration and 6 to immune function. Of the top 20 adhesion-related processes, 11 were involved in the regulation of adhesion, 9 in cell–cell adhesion, and 7 in substrate or matrix interaction. Consistent with GO results showing a dominance of immune-related terms among downregulated processes, half of the top 20 immunity-associated pathways were related to the regulation of immune responses.

### Modulation of BMDC migration and adhesion through CD44

To determine whether the transcriptomic alterations resulted in functional phenotypic changes, we assessed the migration and adhesion abilities of BMDCs. First, isolated BMDCs were tested for their ability to migrate towards CXCL12, CCL2 or CCL5, factors that have been described to be abundant in the pre-metastatic niche^[29]^. The expression of these factors was shown to be increased in FC1245-derived tumors as compared to normal pancreas (Figure 3). Prior to the assay, the BMDCs were activated with CM from a triple culture containing FC1245 cells co-cultured with murine immortalized pancreatic stellate cells (imPSCs) and syngeneic macrophages (Figure 7A). In all cases, migration toward the aforementioned cytokines was reduced upon CD44 inhibition using the anti-CD44 antibody IM7 (Figure 7B). Specifically, for CCL2 and CCL5, chemotaxis was impaired to a similar extent as observed with antibodies targeting their respective receptors, CCR1/2/5 for CCL2 and CCR5/CXCR3 for CCL5. Migration towards CM, comprising a variety of different chemokines, cytokines and growth factors, was drastically reduced upon CD44 inhibition (Supplementary Figure 7A). We have also tested the migration towards CXCL12 of BMDCs isolated from TBM and confirmed a reduced migration upon CD44 blocking (Supplementary Figure 7B). Adhesion to VCAM-1 was blocked upon inhibition of CD44 by IM7, similarly to the treatment with an antibody against integrin α4 (Figure 7C). Moreover, adhesion to fibronectin, a major component of the metastatic niche^[20]^, was completely blocked upon inhibition of CD44 with IM7. In contrast, BMDCs barely adhered to type I collagen, which was not affected by the inhibition of CD44.

**Figure 7.**
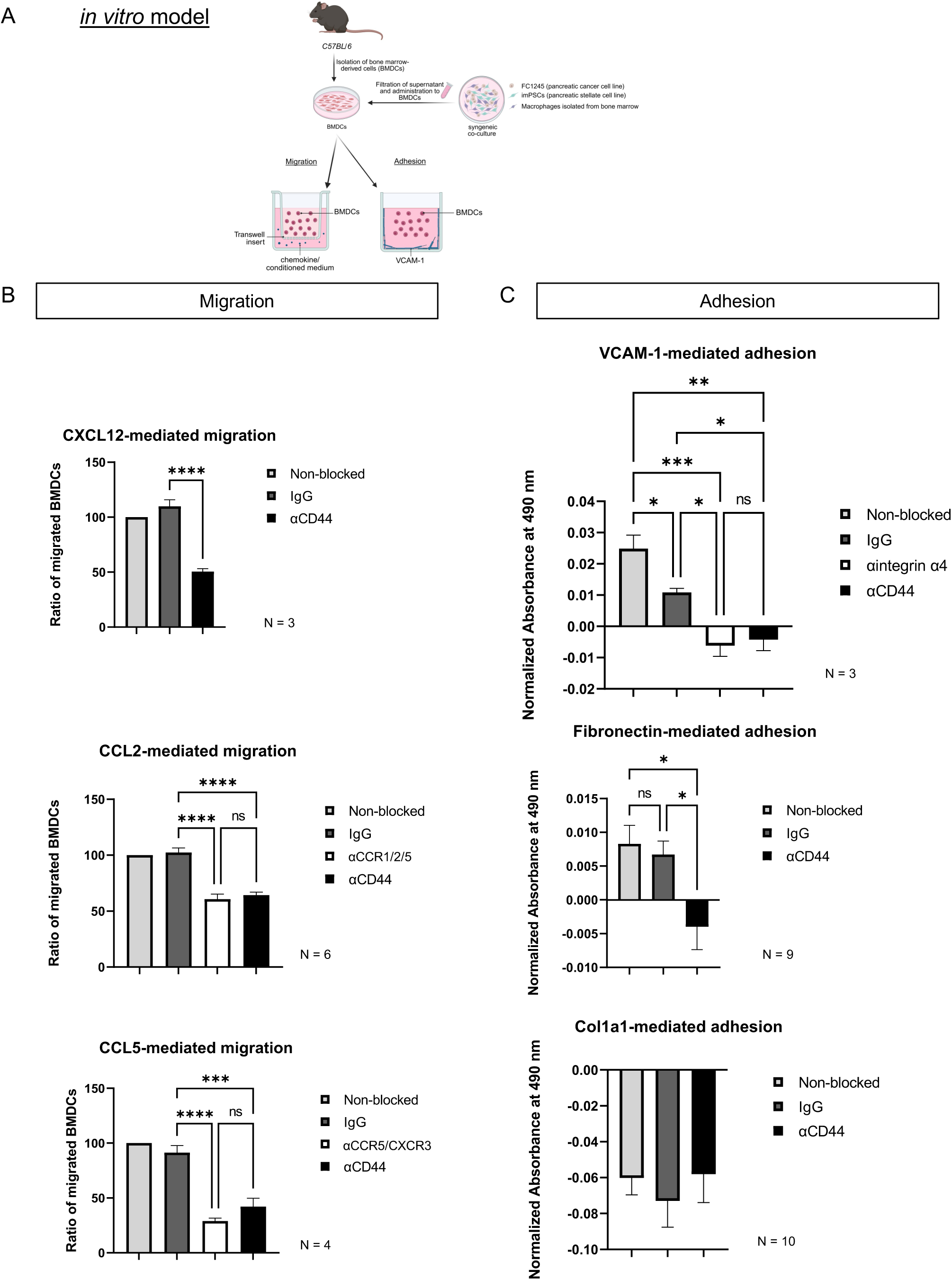
CD44 blockade on BMDCs impairs migration in response to tumor-derived factors and chemokines and adhesion to VCAM-1 and fibronectin but not collagen type I α1. BMDCs were isolated from tibia and femur of healthy C57BL/6 mice. FC1245 PDAC cells were co-cultured with syngeneic immortalized pancreatic stellate cells (imPSC) and bone marrow-derived macrophages (BMDM). The resulting conditioned medium (CM) was applied to BMDCs.**A** Schematic illustration of CM application. Created in BioRender. Mehner, L. (2025) https://BioRender.com/tjbwig4.B BMDCs were cultured for 3 days and subsequently seeded in transwell inserts. BMDCs were allowed to migrate for 5 hours in the presence of chemokines. Migrated cells were counted. BMDCs were treated with inhibitors of CCR1/2/5 (BX471), CXCR4 (AMD3100), a CD44 blocking antibody (clone: IM7), or IgG control. Migration towards CXCL12 was assessed in BMDCs treated with IM7 or IgG control, migration towards CCL2 was assessed in BMDCs treated with BX471, IM7, or IgG control, migration towards CCL5 was assessed in BMDCs treated with TAK779, IM7, or IgG control. **C** CM administered to BMDCs for 7 days. BMDCs were pre-treated for 30 minutes with antibodies targeting integrin α4 (PS/2), CD44 (IM7), or IgG control. Adhesion to VCAM-1 was assessed after 15 minutes, to fibronectin after 25 minutes, and to collagen type I α1 (Col1a1) after 60 minutes. Adhered cells were stained with crystal violet and absorbance was measured at 490 nm using an ELISA plate reader. Schematic illustration was created using BioRender.com. Migration: Statistical significance was determined using the one-sided unpaired Student’s t test for CXCL12-mediated migration and ordinary one-way analysis of variances (ANOVA) and Holm-Šídák’s multiple comparisons post-hoc test for remaining analysis; ns=not significant, *p-value<0.0332; **p-value<0.0021; ***p-value<0.00021; ****p-value<0,0001. Adhesion: Statistical significance was determined using the ordinary one-way analysis of variances (ANOVA) and Holm-Šídák’s multiple comparisons post-hoc test; ns=not significant, *p-value<0.0332; **p-value<0.0021; ***p-value<0.00021; ****p-value<0,0001.

### CD44 modulates BMDC inflammatory and immune function

The RNAseq data indicated that CD44 may play a role in regulating the immune response (Figure 6F). To investigate whether *Cd44* removal affects the immunosuppressive properties of BMDCs, we measured the expression of several key immunosuppressive factors in BMDCs of *Cd44^ΔHem^* mice compared to *Cd44^+/+^*mice. The expression of Arginase-1 (Arg1), involved in the depletion of arginine, an amino acid that is essential for the formation of the T cell receptor and of the TCR-ζ chain in particular, was measured by qPCR (Figure 8A) and in ELISA (Figure 8C)^[30]^. At the mRNA and protein levels, it was decreased in BMDCs isolated from *Cd44^ΔHem^* mice. The same observation was made in the case of *Ido1*, leading to depletion of the essential amino acid tryptophane. This enzyme was decreased in BMDCs isolated from *Cd44^ΔHem^* mice. IL-10, involved in the generation of regulatory T cells (T_Reg_) cells and exerting direct immunosuppressive function on T effector cells, was also decreased in BMDCs isolated from *Cd44^ΔHem^* mice. IFNβ1 and IFNγ, which can facilitate T_Reg_ phenotypes and suppress T cell activation, were additionally decreased. Upon abrogation of *Cd44*, mRNA levels of *iNos* are increased. iNOS is commonly expressed by M1 macrophages in response to acute inflammation. PD-L1, which negatively regulates T cell function and facilitates immune evasion by tumor cells, was found to be downregulated (Figure 8B). Also, the expression of the receptor of PD-L1, PD-1 decreased. PD-1 is also present on T cells suggesting that decreased expression of PD-L1, coherent with findings of other groups identifying CD44 as a key regulator of PD-L1 expression in non-small lung cell carcinoma and triple-negative breast cancer^[31]^. Hence, BMDCs might be less immunosuppressive in the absence of CD44.

**Figure 8.**
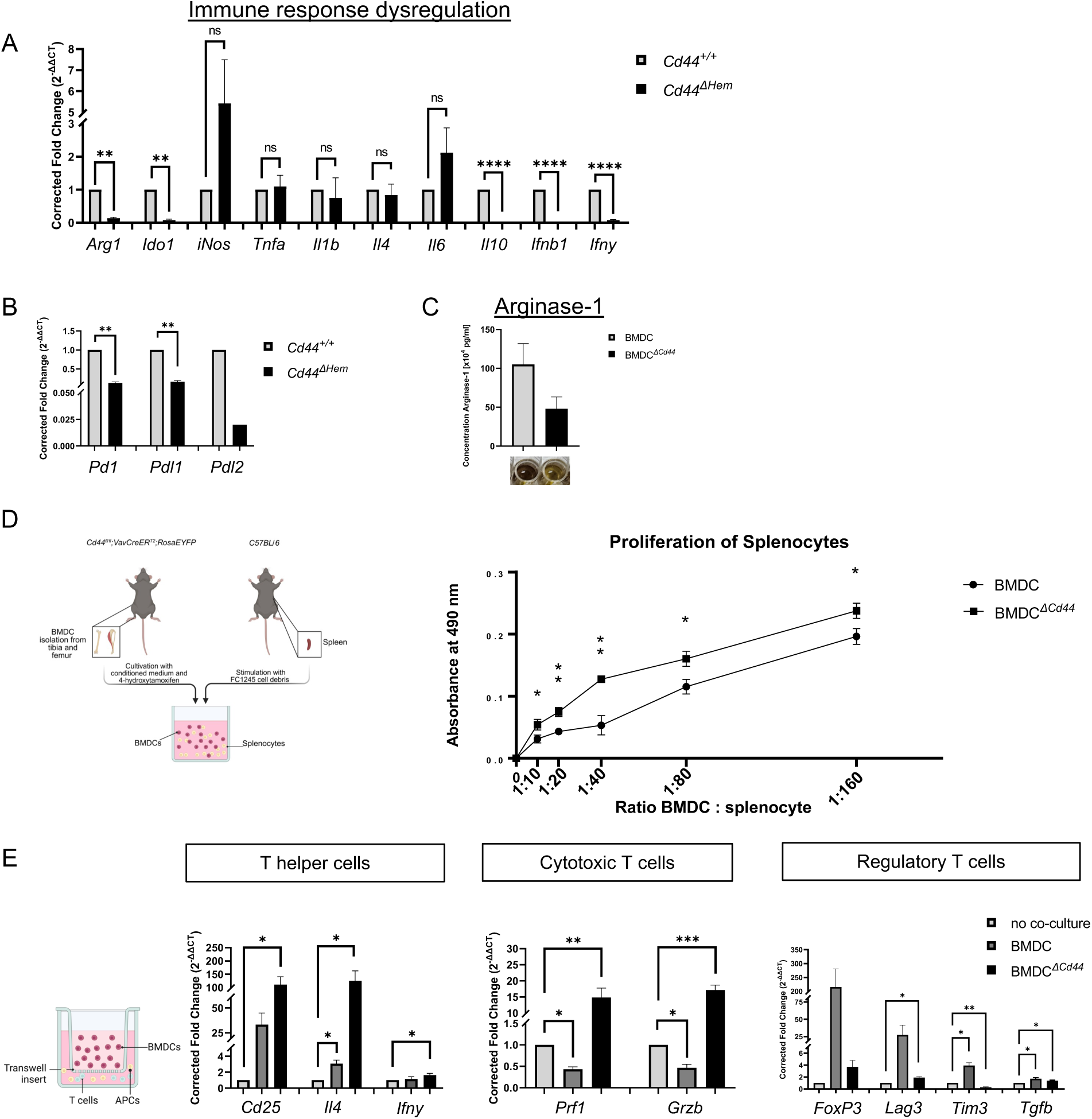
CD44 abrogation on BMDCs reduces immunosuppressive potential in the context of PDAC. ***In vivo* setup:** *Cd44^fl/fl^;VavCreER^T2^;RosaEYFP* (*Cd44^ΔHem^*) and control (*Cd44^+/+^*) animals were injected with tamoxifen for five consecutive days. FC1245 PDAC cells were injected into the pancreas at day 0, and animals were sacrificed at 14 or 21 dpi for BMDC isolation from tibia and femur. ***In vitro* setup**: BMDCs were isolated from *Cd44^+/+^*and *Cd44^ΔHem^* animals and cultured with CM from FC1245 PDAC cells, immortalized pancreatic stellate cells (imPSCs) and BMDMs. **A**,**B** mRNA levels of immune-related genes were analyzed by qPCR (normalized to *Cd44^+/+^*). **C** Quantification of Arginase-1 by ELISA from BMDC lysates. **D** Syngeneic splenocytes were cultured and treated with FC1245 cell debris. 24 hours post treatment, BMDCs and BMDC*^ΔCd44^* were co-cultured with splenocytes at varying ratios (1:10, 1:20, 1:40, 1:80, 1:160). After 24 hours, metabolic activity was analyzed by conversion of a tetrazolium compound into formazan and the resulting change in absorbance (CellTitre96 AQueous One Solution Reagent, measured by SpectraMax). Schematic Illustration: Created in BioRender. Mehner, L. (2025) https://BioRender.com/r3jw4fi **E** Expression level of mRNA of immune-related genes was analyzed by qPCR (normalized to *Cd44^+/+^*). Schematic Illustration: Created in BioRender. Mehner, L. (2025) https://BioRender.com/k0eawdc

To directly assess the immunosuppressive capability of BMDCs, they were incubated with CM from the triple co-culture and incubated with splenocytes that had been exposed to FC1245 debris (Figure 8D). Proliferation of splenocytes co-cultured with BMDC*^ΔCd44^* was increased compared to splenocytes co-cultured with control BMDCs. This result confirms the aforementioned data, suggesting that CD44 confers an immunosuppressive character to BMDCs. In another context, splenocytes isolated from *B6.Cg-Tg(TcraTcrb)425Cbn/J* animals (OT-II mice) were treated with ovalbumin and co-cultured with BMDCs expressing CD44 or not in a boyden chamber. Measurement of mRNA levels of markers of T helper cells revealed an increase of *Cd25* and *Il4* in the absence of CD44 on BMDCs (Figure 8E). Although the expression of *Ifng* was low, it was increased in the context of incubation with BMDC*^ΔCd44^* cells. Markers of cytotoxic cells such as *perforin 1* and *granzyme B* were induced when splenocytes were indirectly co-cultured with BMDC*^ΔCd44^*. Markers of T_Reg_ cells were decreased in particular *Lag3* and *Tim3*.

We have additionally tested TGF-β-mediated signaling in BMDCs from *Cd44^+/+^* and *Cd44^ΔHem^*. Not only did CD44 form a complex with TGFβRI (Supplementary figure 8A), it also influenced its downstream signaling as seen for SMAD3 phosphorylation and *Garp* expression (Supplementary Figure 8B and C). Bulk RNAseq further revealed an involvement of CD44 in TGF-β receptor activation (Supplementary Figure 8D) Additionally, genes related to the IL-6/JAK/STAT3 pathway were enriched in BMDCs of TBM compared to NTBM, as demonstrated by the running enrichment score (Supplementary 8E). Strikingly, upon the knockout of *Cd44* on BMDCs, enrichment decreased.

Similar experiments were performed with BMDCs isolated from *Cd44v6^ΔHem^*mice compared to *Cd44v6^+/+^*mice (Figure 9). The same factors (*Arg1*, *Ido1* and *Ifnb1*) were found to be decreased upon removal of Cd44v6 in BMDCs from tumor-bearing mice (Figure 9A). On the protein level Arg1 was clearly downregulated (Figure 9C). In contrast *iNos* was increased (Figure 9A).

**Figure 9.**
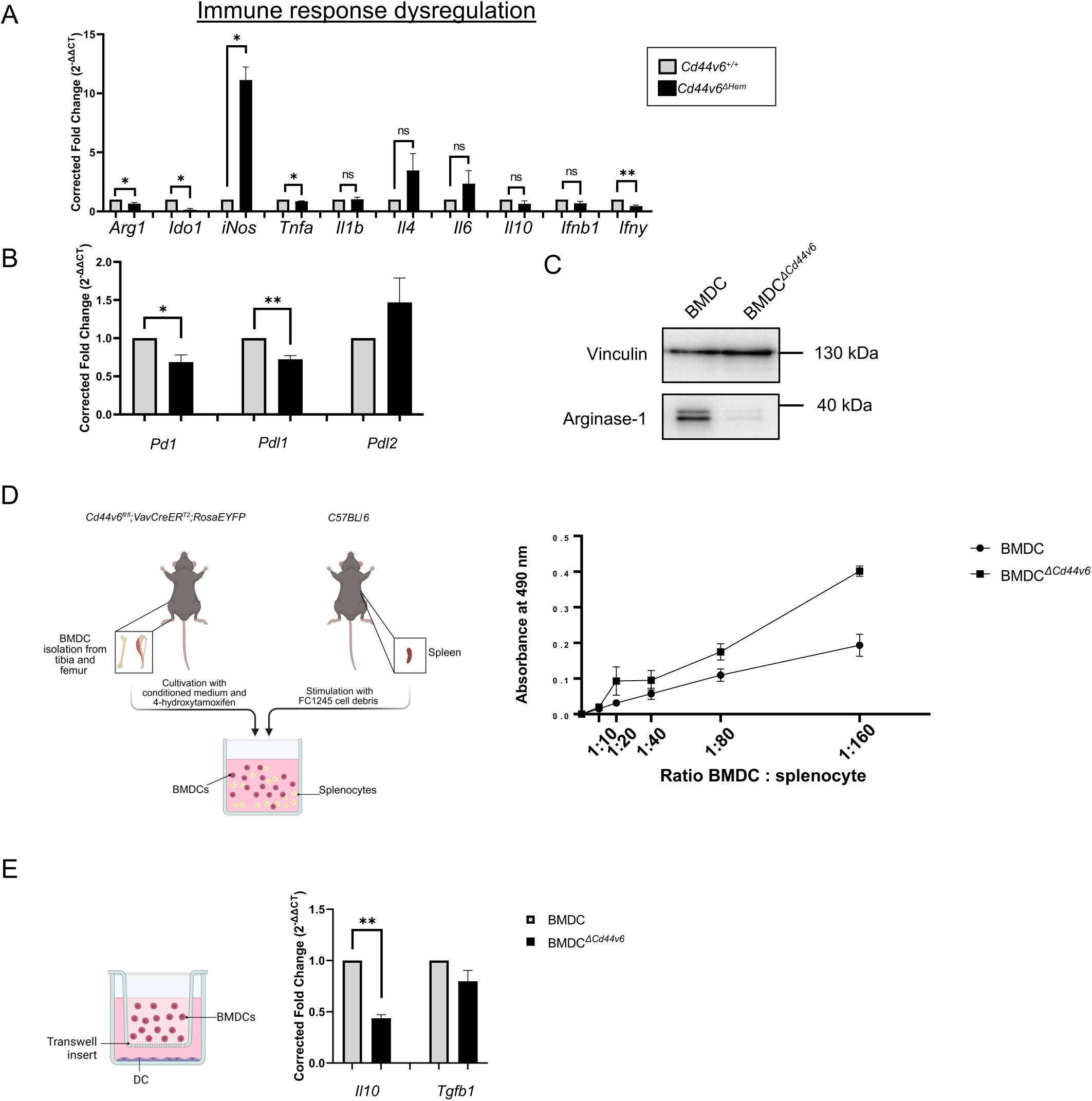
Knockout of the *Cd44v6* isoform on BMDCs decreases immunosuppressive potential. **A,B** *Cd44v6^ΔHem^* and control *Cd44v6^+/+^* animals were injected with tamoxifen intraperitoneally for five consecutive days. FC1245 PDAC cells were injected orthotopically, and mice were sacrificed at 14 dpi. BMDCs were isolated from tibia and femur, and RNA was extracted for analysis of the expression patterns of **A** immunity-related mediators and **B** immune checkpoint receptors and ligands. Corrected fold change (2^−ΔΔCT^) was calculated by normalizing *Cd44v6^ΔHem^ to Cd44v6^+/+^*. Gapdh was used as a reference gene. **C** Protein lysates of BMDCs and BMDC*^ΔCd44v6^* were subjected to western blot analysis for Arginase-1, with vinculin as a loading control. **D** Splenocytes from syngeneic animals were cultured and treated with FC1245 cell debris. After 24 hours of treatment, BMDCs and BMDC*^ΔCd44v6^* were co-cultured with splenocytes at varying ratios (1:10, 1:20, 1:40, 1:80, 1:160). Metabolic activity was analyzed 24 hours later by measuring absorbance after conversion of a tetrazolium compound into formazan (CellTitre96 AQueous One Solution Reagent, measured by SpectraMax). Schematic Illustration: Created in BioRender. Mehner, L. (2025) https://BioRender.com/puo4w50 **E** BMDCs and BMDC*^ΔCd44v6^*were co-cultured with the dendritic cell line DC2.4 in a transwell system for 5 days. Expression patterns of DCs were analyzed by qPCR. Corrected fold change (2^−ΔΔCT^) for dendritic cells co-cultured with BMDC*^ΔCd44v6^* was calculated and normalized to DC cultured with BMDCs. *Gapdh* was used as a reference gene. Schematic Illustration: Created in BioRender. Mehner, L. (2025) https://BioRender.com/3bcue4c

In the case of immune checkpoint inhibitors, expression of *Pdl2* was unchanged whereas *Pd1* and *Pdl1* were decreased (Figure 9B). A proliferation assay was performed using splenocytes of healthy mice incubated with BMDCs expressing CD44v6 or not. Again, splenocytes incubated with BMDC*^ΔCd44v6^* cells proliferated more (Figure 9D). BMDC*^ΔCd44v6^*failed to induce expression of immunosuppressive IL-10 and TGF-β1 in indirect co-culture with DCs (Figure 9E).

## Discussion

Using immunocompetent mice orthotopically injected with PDAC cells, we show that BMDCs establish an immunosuppressive milieu which promotes the establishment of cancer cells in the liver. Clusters of BMDCs can be found in the liver that highly express CD44 and its variant isoform CD44v6. BMDC mobilization and homing in pancreatic cancer depend on CD44. Selective deletion of *Cd44/Cd44v6* in BMDCs significantly disrupts niche formation in the liver and reduces metastatic burden. Thus, CD44 emerges as a promising target to modulate metastatic niche development. Through an unbiased approach, we found that *Cd44* loss impairs BMDC mobilization and immunosuppressive functions. These insights highlighting distinct CD44-dependent mechanisms driving metastasis when expressed on cancer cells or on BMDCs^[16]^ (Figure 10).

CD45^+^ cell clusters observed in the livers of TBM are recruited from the bone marrow, as some of these cells expressed markers of hematopoietic stem cells (Sca1^+^) and progenitor cells (c-kit^+^), while the cells scattered around the clusters in the liver did not. The general role of c-kit⁺/Sca1⁺ cell mobilization to the liver in pathological contexts remains unclear, though they may aid regeneration after liver injury^[32]^. A microenvironment expressing regenerative stimuli may aid in the survival and development of incoming disseminated cancer cells increasing the likelihood to prosper into metastases. Furthermore, the transcriptional changes in cells isolated from bone marrow of TBM, compared to bone marrow cells from NTBM argues in favor of a reaction to stimuli generated by the primary tumor directly or indirectly through stimuli induced in the liver by tumor-derived factors^[5, 33, 34]^. Among the significantly upregulated genes in BMDCs of TBM, is *Cd44*. Interestingly, the expression of CD44 on CD34^+^ hematopoietic progenitor cells has been found to be significantly lower when found in the bone marrow than in systemic circulation^[35]^. Myeloid progenitors are mainly found in the CD34^+^CD44^+^ cell fraction. Additionally, the *Cd44* transcripts in BMDCs of TBMs, exhibit a splicing skew towards a significant inclusion of CD44 variant exons. This observation underscores CD44 as a molecular target in these cells. Beyond its high expression, CD44 offers the opportunity to target only isoforms specifically associated with the metastatic process sparing isoforms involved in physiological tasks in other tissues. This has the potential to reduce the toxicity of agents that target these specific isoforms. In the present work the genetic loss of function of *Cd44v6* in BMDCs had similar effects to those accomplished by full *Cd44* knockout.

To understand the biological processes affected by the *Cd44* knockout on BMDCs we undertook and unbiased approach through RNAseq Gene Ontology (GO) analysis. Since GO analysis showed that the knockout of *Cd44* affected migration, adhesion, and immune response processes, we devised an *in vitro* system to functionally test the consequences of the transcriptional changes observed. BMDCs were exposed to factors of a CM coming from a triple co-culture system including FC1245 cells, stellate cells and macrophages, two cell types belonging to the PDAC stroma. CD44 blocking by a specific antibody, indeed affected cell migration in trans-well assays towards different chemokines, at least to the same extent that specific chemokine receptor inhibitors achieved. In the same way CD44 blocking prevented BMDC adhesion to wells coated with VCAM-1, involved in rolling of circulating cell before extravasation, and extracellular matrix proteins^[36, 37]^. These aspects could affect BMDC homing and extravasation to form cell clusters in the liver, compatible with the stark decrease of cells cluster formation in tumor bearing mice with a *Cd44* knockout. Of note, each one of these assays involved different chemokine receptors and integrins of which their function could be impaired by the blocking of CD44. We have proven the physical interaction and cooperation of CD44 with the function of CXCR4^[38][2]^. In a similar way, CD44 has been proven to enhance the function of integrins α4 and β1 (VLA-4)^[39–41]^. However, the effect on the function of other chemokine receptors and integrins in these assays, argues in favor of an even broader pleiotropic function of Cd44, and underscores the notion of molecular target to modulates diverse cellular functions.

The vav promoter has been reported to be expressed in all hematopoietic lineages^[27]^. The *Cd44/Cd44v6* knockout model significantly reduced the number of CD45^+^ cell clusters in the liver of tumor bearing mice at 14 dpi to approximately 30% compared to controls, as well as metastasis formation at 21 dpi. In our mechanistic experiments *in vitro*, this knockout prevented the immunosuppressive stimuli of BMDCs on splenocytes. Interestingly, in our mechanistic *in vitro* experiments, blocking of CD44 with a specific antibody, had a very steep effect on BMDC migration and adhesion. The significant effect of a loss-of-function and that may be achieved or even improved by using blocking molecules, underscores the translational potential of these findings.

The overall 5-year survival of patients with PDAC even localized in stage IA treated with curative intent through surgery and adjuvant chemotherapy is of only 50%, showing that PDAC undergoes an early dissemination^[2, 42]^. At 14 dpi of KPC (FC1245) cells in our syngeneic PDAC model, we observed CD45⁺ hematopoietic cell clusters in the liver, along with a few scattered Pdx1⁺ pancreatic cancer cells. One week later, at 21 dpi, we detected micrometastases in which Pdx1^+^ and CD45^+^ cells coexisted to a similar extent. These findings support a model in which hematopoietic cells remain key components of the metastatic niche throughout metastasis progression. Studying these cell clusters after initial cancer cell seeding and even after micrometastases have formed remains highly relevant, because metastatic lethality is driven by overall tumor burden, not merely initial spread. Given evidence that cells from established metastases can seed new ones, targeting this branching progression could help contain existing lesions and prevent further spread, aligning more closely with clinical realities^[43]^.

The mechanisms of how CD44 may affect immune functions are complex. In this work we devised several assays to investigate this phenomenon. From a gene expression perspective, CD44 and CD44v6 were involved in the expression of immunosuppressive mediators *Arg1*, *Ido1*, interleukins and interferons. From a functional point of view, we demonstrated that splenocyte proliferation was increased upon loss of *Cd44* or *Cd44v6*. Additionally, BMDCs expressing CD44 favored a regulatory phenotype in T cells, and CD44v6^+^ BMDCs favored a regulatory phenotype in DCs. We are suspecting that the immunosuppressive characteristic of CD44 can be ascribed to CD44v6, and that its expression promotes a myeloid-derived suppressor phenotype in BMDCs. Targeting CD44v6 *in vivo* might therefore reprogram immunosuppressive cells and promote immune detection of incoming cancer cells, drastically decreasing metastatic burden.

## Methods

### Mice

*C57BL/6-Tg(VavCreER^T2^)* mice were crossed with *C57BL/6-Tg(R26-LSL-EYFP^KI/KI^)* to generate the *C57BL/6-Tg(VavCreER^T2^/STOP-YFP*. C57BL/6-Tg(VavCreERT2) mice were kindly provided by the Walter and Eliza Hall Institute of Medical Research (WEHI) and Dr. Marco Harold. This resulting mouse line was subsequently crossed with *C57BL/6-Tg(Cd44-[exon3]^flox/flox^*mice to generate the *Cd44^flox/flox^Vav-CreER^T2^/STOP-EYFP* mouse line, called *ΔHem* for brevity. The *C57BL/6-Tg(VavCreER^T2^/STOP-YFP)* was also crossed with the *C57BL/6-Tg(Cd44v6^flox/flox^)* to generate the *Cd44v6^flox/flox^Vav-CreER^T2^/STOP-EYFP* mouse line. The resulting lines were used for *Cd44* or *Cd44v6* bone marrow-derived cell knockout experiments respectively. The animals were housed and maintained under specific pathogen-free conditions in facilities approved by the Regierungspräsidium Karlsruhe. All animals were handled according to EU directives for animal experimentation. The experiments were authorized by the Regierungspräsidium (35-9185.81/G-158/18 and 35-9185.81/G-286/15).

### Cell lines

FC1245 cells were a kind gift from Dr. Dave Tuveson (Cold Spring Harbor Laboratory, Cold Spring Harbor, NY), and were cultured in DMEM GlutaMAX supplement (Gibco, MA, USA) and 10% fetal calf serum (FCS). The FC1245 cell line derives from a KPC mouse containing a point mutation in TP53 gene (R172H), and a point mutation in the KRAS gene (G12D). imPSCs were obtained from Dr. Angela Mathison and Dr. Raul Urrutia (Medical College of Wisconsin, Milwaukee, USA)^[44]^. They were cultured in DMEM GlutaMAX supplement (Gibco, MA, USA) and 10% fetal calf serum (FCS).

### Orthotopic PDAC cell implantation

Eight to twelve week-old *Cd44^ΔHem^* mice or controls, were injected intraperitoneally with 100µL of Tamoxifen diluted in peanut oil at a concentration of 20 mg/ml, once a day, for 5 days. At the seventh day after the initiation of Tamoxifen injection, 1 × 10^5^/ml FC1245 cells were suspended in sterile PBS and put on ice. At the time of injection, the cells were resuspended by inverting the tube several times. A 1-ml syringe with a 27G of ½ -inch needle and left on ice. The animals were anesthetized by isofluorane inhalation vaporized with compressed air at 1 liter per minute in supine position, without fixation. The abdominal skin was cleaned with 70% ethanol. A 0.5 cm incision was made with sterile micro-scissors at the left abdominal flank. The spleen and part of the pancreas was exposed out through the incision by gently pulling with blunt forceps. 30 µL of cell suspension were injected into the pancreas, 3 mm medial to the spleen hilium. The pancreas was kept externalized and untouched for 1 minute to inspect for hemorrhage and leakage. The pancreas and spleen were returned to the peritoneal cavity with blunt forceps, and muscle and skin layers were sequentially close with interrupted 4-0 non-reabsorbable sutures.

### BMDC harvesting

Bone marrow-derived cells were obtained by harvesting the bone marrow from the tibia and femur of mice. The bone marrow was then put through a 35 µm filter, washed with ammonium-chloride-potassium (ACK) to lyse red blood cells, resuspended in PBS, centrifuged for 5 minutes at 1400 RPM, resuspended in IMDM medium with 30% FSC and cultured at 37°C.

### Flow cytometry

Mouse bone marrow-derived cells, freshly harvested and cultured, were treated with 4-OH-tamoxifen (4OHT) 1µM (Sigma) for 7 days. They were harvested by cell scraping and incubated with mouse Fc Block^TM^ (Purified rat anti-mouse CD16/CD32; BD PharmingenTM) for 30 minutes, then stained using a human/mouse anti-CD44 antibody (IM7), directly coupled to phycoerythrin (PE) (Biolegend) or isotype control, for 1 hour. They were washed 3x with blocking buffer (PBS + 5%FBS) and analyzed by flow cytometry using a FACSAria Fusion cytometer (BD).

### Bulk RNASequencing

Paired-end FASTQ files were processed with Trimmomatic (v0.39) to remove adapters and low-quality reads. High-quality reads were aligned to the mouse genome (mm10) using STAR (v2.7.11a), and gene counts were quantified. Downstream analysis in R (v4.4.0) included read count normalization and differential expression analysis with DESeq2. Gene-set enrichment analysis (GSEA) was performed using ClusterProfiler with MSigDB as the reference. Significance was determined using a p-value threshold of 0.05.

### Hematoxylin and Eosin staining

FFPE tissue sections were deparaffinized in xylene (2 × 5 min), rehydrated through graded ethanol (100%, 95%, 70%; 1 min each), and rinsed in tap water (2 min). Sections were stained with hematoxylin (1 min), washed in tap water (5 min), and counterstained with eosin (1 min; 0.1% working solution in 1% acetic acid, 70% ethanol). Slides were dehydrated through ethanol (70%, 95%, 100%; 2 × 1 min each), cleared in xylene (2 × 1 min), and mounted with Eukitt. Metastases were assessed by Dr. Matthias Gaida (University Medical Center Mainz/TRON), who also provided representative images.

### Direct Immunofluorescence

Paraffin-embedded tissue sections were prepared as described for IHC up to the peroxidase blocking step. Sections were treated with a blocking solution containing 5% FBS in PBS-T for 1 hour at room temperature and then incubated with the antibody directly coupled to a fluorophore or the at room temperature for 2 hours in blocking buffer, were washed in PBS-T, mounted with fluorescence mounting medium (Dako). Micrographs were taken using a Zeiss LSM 800 confocal microscope. Images were processed using Fiji (U.S. NIH).

### RNA extraction and reverse transcription and qPCR

Total RNA was extracted combining Trizol (Invitrogen) and the RNeasy Mini Kit (Qiagen) according to the protocol by A. Untergasser (www.untergasser.de/lab). Prior to reverse transcription for cDNA synthesis, DNase I was added to the isolated RNA (30 min at 37 °C). Next, DNase stop solution (Promega) containing EDTA (10 min at 65 °C) and random primers were added (1 µl, 5 min at 70 °C). cDNA synthesis reactions (10 µl) were prepared of 4 µL 5× MLVRT buffer (Promega), 2 µl dNTPs and 1 µl reverse transcriptase (MLVRT; Promega). Samples were incubated 10 min at 25 °C followed by 60 min incubation at 42 °C. The qPCR reactions (20 µl) consisted of 10 µl 2× GoTaq® qPCR Master Mix (Promega), 0.5 mM reverse and forward primers, and 2 µl cDNA. The qPCR was performed according on the CFX96 Touch Real-Time PCR detection system (Biorad). *Gapdh* and *β-Actin* were used as reference genes for relative quantification according to the Litvak method (ΔΔCt). Primers were designed using the Primer3 open software (http://primer3.ut.ee) based on NCBI-database transcript sequences, and are detailed in Supplementary Table 1.

### Migration

BMDCs were isolated from tibia and femur of C57BL/6 mice and cultured for 3 days in syngeneic CM. After 1 h serum starvation, cells were collected, counted (2 × 10⁶ cells/ml), and pre-treated with Fc block (1 µl/10⁶ cells) for 30 min, followed by CD44 antibody (IM7, 100 µg/ml), CXCR4 inhibitor (AMD3100), CCR1/2/5 inhibitor (BX471), CXCR3/CCR5 inhibitor (TAK779), or isotype control.

For migration, 2 × 10⁶ BMDCs in 1 ml were seeded in 6-well ThinCert inserts (8 µm pores); for CCL5 assays, 2 × 10⁵ cells in 100 µl were seeded in 12-well inserts. The lower chambers contained either CM or chemokines (CCL2: 200 ng/ml, CXCL12: 50 ng/ml, or CCL5: 100 ng/ml) in IMDM with 1% FCS. After 5 h at 37 °C, migrated cells were collected, washed, and quantified using an Eve automated cell counter. Migration was normalized to non-blocked controls.

### Adhesion

BMDCs were isolated from C57BL/6 mice and cultured for 7 days in syngeneic CM. On day 6, 96-well plates were coated overnight at 4 °C with VCAM-1 (10 µg/ml), fibronectin (25 µg/ml), COL1A1 (50 µg/ml), or diluent controls. Wells were washed with PBS on day 7. BMDCs were starved for 1 h, detached with Accutase, and adjusted to 2.5 × 10⁶ cells/ml. Cells were Fc-blocked (1 µl/10⁶ cells, 30 min), then treated with CD44 (IM7, 100 µg/ml), integrin α4/CD49d (PS/2, 2 µg/10⁶ cells), or IgG control antibodies. After 30 min, 2.5 × 10⁵ cells/well were seeded in triplicates. Plates were centrifuged (200 rpm, 1 min) and incubated (37 °C, 5% CO₂) for 15–60 min depending on the coating. Wells were washed with ice-cold PBS (+Ca²⁺/Mg²⁺), fixed with −20 °C methanol, and stained with 0.5% crystal violet. Dye was solubilized with methanol, and absorbance was measured at 590 nm. Adhesion was normalized to non-blocked controls, and diluent-coated background was subtracted.

### Co-cultures

BMDCs were isolated from *Cd44^ΔHem^* or *Cd44v6^ΔHem^* mice and respective controls, and cultured for ≥6 days in syngeneic CM with daily 4OHT (1 μM).

#### BMDC and Splenocyte Co-culture

On day 6, splenocytes from the same strains were isolated and cultured in RPMI supplemented with 10% FCS, 1% PenStrep, non-essential amino acids, 500 μM HEPES, and 50 μM 2-mercaptoethanol. On day 7, splenocytes were stimulated with FC1245 cell debris (40 μl/ml for 24 h). On day 8, BMDCs and splenocytes were co-cultured at ratios of 1:0, 1:10, 1:20, 1:40, 1:80, and 1:160 in triplicates (BMDCs at 1 × 10⁵/ml; splenocytes at 1–16 × 10⁶/ml). After 12 h, 20 μl of FC1245 debris was added per well. On day 9, cell viability was assessed using Promega’s CellTiter 96 AQueous One Solution (20 μl/well, 2 h incubation), and absorbance was measured at 490 nm. Values were normalized to BMDC-only controls.

#### Indirect Co-culture with OT-II T Cells

BMDCs from *Cd44^ΔHem^* mice were cultured with daily 4OHT in CM for 6 days. Splenocytes from OT-II mice were cultured as above and pulsed with OVA^323–339^ peptide (250 ng/ml) on day 3. On day 6, 1 × 10⁵ BMDCs were seeded in ThinCert inserts (0.4 μm pore) in 12-well plates; 1 × 10⁶ OT-II splenocytes were added to the lower chamber. IL-2 (1 ng/ml) was added on day 7. On day 10, RNA was extracted from splenocytes for downstream analysis.

Indirect Co-culture with DC2.4 Cells

BMDCs from *Cd44v6^ΔHem^* mice were cultured for 7 days with daily 4OHT. On day 8, 2 × 10⁵ BMDCs were seeded in ThinCert inserts (0.4 μm pores) and co-cultured with 5 × 10⁵ DC2.4 cells in 6-well plates. After 48 h, RNA was extracted from DC2.4 cells for transcriptomic analysis.

### Protein isolation and Protein detection

Cells were lysed with 1× RIPA buffer. Lysates were collected using a rubber scraper and sonicated on ice for 15 min. After centrifugation (13,200 rpm, 20 min, 4 °C), the supernatant was transferred to fresh tubes. Laemmli sample buffer (4×) with 10% β-mercaptoethanol was added, and samples were boiled at 95 °C for 5 min. Lysates were stored at −20 °C until SDS-PAGE.

#### IP

BMDCs were lysed using either 1% Triton X-100 buffer. A 50 μl aliquot of lysate was reserved as input, mixed with 4× Laemmli buffer, and stored at −20 °C. Lysates were pre-cleared with 30 μl Protein G agarose beads (washed 3× with lysis buffer, 1,200 rpm, 3 min). After centrifugation, pre-cleared lysates were incubated overnight at 4 °C with anti-CD44 (KM201, 5 μg/200 μl lysate) or anti-CXCR4 (3 μg/200 μl lysate), or matched IgG controls. After antibody incubation, 30 μl of washed Protein G agarose beads were added and rotated for 2 h at 4 °C. Beads were washed 3×with lysis buffer, resuspended in 50 μl 2× Laemmli buffer with 10% β-mercaptoethanol, boiled at 95 °C for 5 min, and stored at −20 °C. Co-immunoprecipitated proteins were analyzed by western blot.

#### Western Blot

Proteins were separated by SDS–PAGE using 5% stacking and 8% separating gels. Electrophoresis was performed at 100 V for 60 min followed by 120 V for 45 min. Proteins were transferred to PVDF membranes using the Trans-Blot Turbo system (Bio-Rad). Membranes were blocked with 5% BSA in TBS-T for 1 h, incubated overnight at 4 °C with primary antibodies (1:1000 in 5% BSA/TBS-T), and washed. HRP-conjugated secondary antibodies (1:2000 in TBS-T) were applied for 1 h at room temperature. After washing, signals were developed using ECL substrate and visualized on a ChemiDoc Touch Imaging System (Bio-Rad).

#### Arginase-1 Elisa

Arginase-1 levels in BMDCs from tumor-bearing mice were quantified using the Mouse Arginase-1 SimpleStep ELISA® Kit (Abcam) according to the manufacturer’s instructions. BMDCs (1 × 10⁶) were lysed in 500 μl of the provided lysis buffer and diluted 1:50 prior to analysis. Sample concentrations were determined by interpolating absorbance values from a standard curve and adjusted for the dilution factor.

### Statistical analysis

Statistical analysis was performed using GraphPad Prism 9.0.0 (GraphPad, RRID:SCR_002798). The tests for statistical inference used for each experiment are indicated in the figure legends. Mean values of quantitative variables between two independent groups were compared using a Student’s t test if datasets had a normal distribution according to Shapiro–Wilk test, or using Mann–Whitney U-test for non-normal distributions. We accepted an α = 0.05 for significant differences.

## Acknowledgements

LMM, LMS, SMT, PH and V.O-R. were supported by the Helmholtz program “Materials Systems Engineering (MSE)”. We thank the Deutsche Forschungsgemeinschaft for the support (OR124/15-1) We acknowledge funding from the German Federal Ministry of Education and Research (BMBF) within the Medical Informatics Funding Scheme EkoEstMed–FKZ 01ZZ2015 (G.A.). We thank Michelle Christ and Aleyna Yildirim for technical help. We thank Dr. Dave Tuveson (Cold Spring Harbor Laboratory, Cold Spring Harbor, NY) for providing the FC1245 cell line. We would also like to thank Dr. Angela Mathison and Dr. Raul Urrutia (Medical College of Wisconsin, Milwaukee, USA) for providing the imPSCs cell line.^[44]^ We thank the Walter and Eliza Hall Institute of Medical Research (WEHI) and Dr. Marco Herold for providing VavCreER^T2^ mice.^[28]^

## Data availability

Data will be made available in a repository upon acceptance.

---

**Scheme 1.**
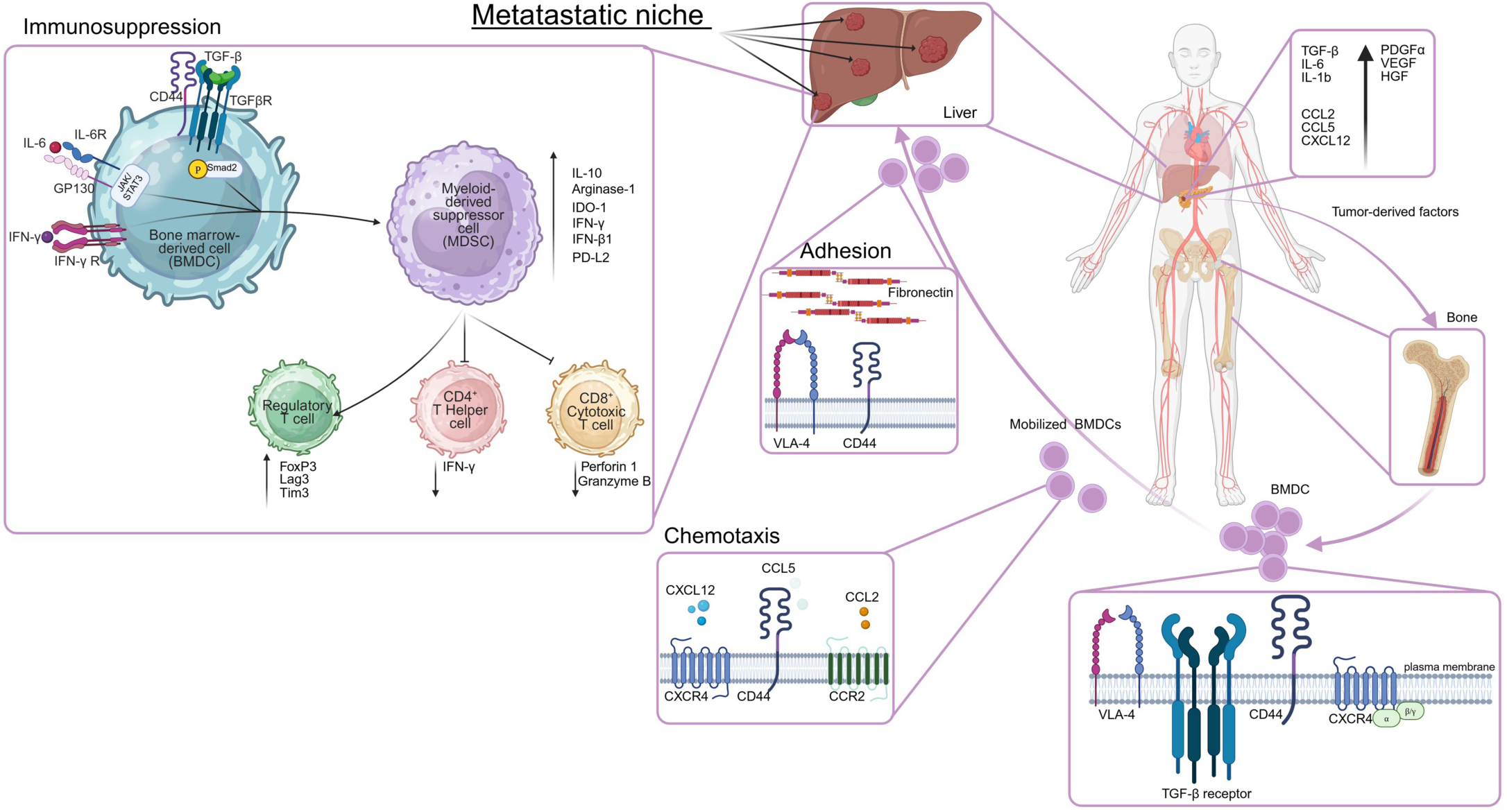
CD44 is critical for the mobilization, migration, and adhesion of bone marrow-derived cells (BMDCs) to metastatic sites in response to tumor-derived signals. It mediates chemokine-driven migration (CCL2, CCL5, CXCL12) and enhances adhesion through VCAM-1 and fibronectin interactions, especially in the fibronectin-rich liver microenvironment induced by PDAC. CD44 and its variant CD44v6 also promote the differentiation of BMDCs into immunosuppressive myeloid-derived suppressor cells (MDSCs) via TGF-β, IL-6/STAT3, and IFN-γ pathways, supporting immune evasion and metastatic niche formation. Created in BioRender. Mehner, L. (2025) https://BioRender.com/bvi5noc

