## Supplementary Data for "Reprogramming Immunosuppressive Bone Marrow–Derived Cells via CD44 Targeting Impacts Pancreatic Cancer Metastasis"

PDAC tumor-bearing animals versus non-tumor-bearing

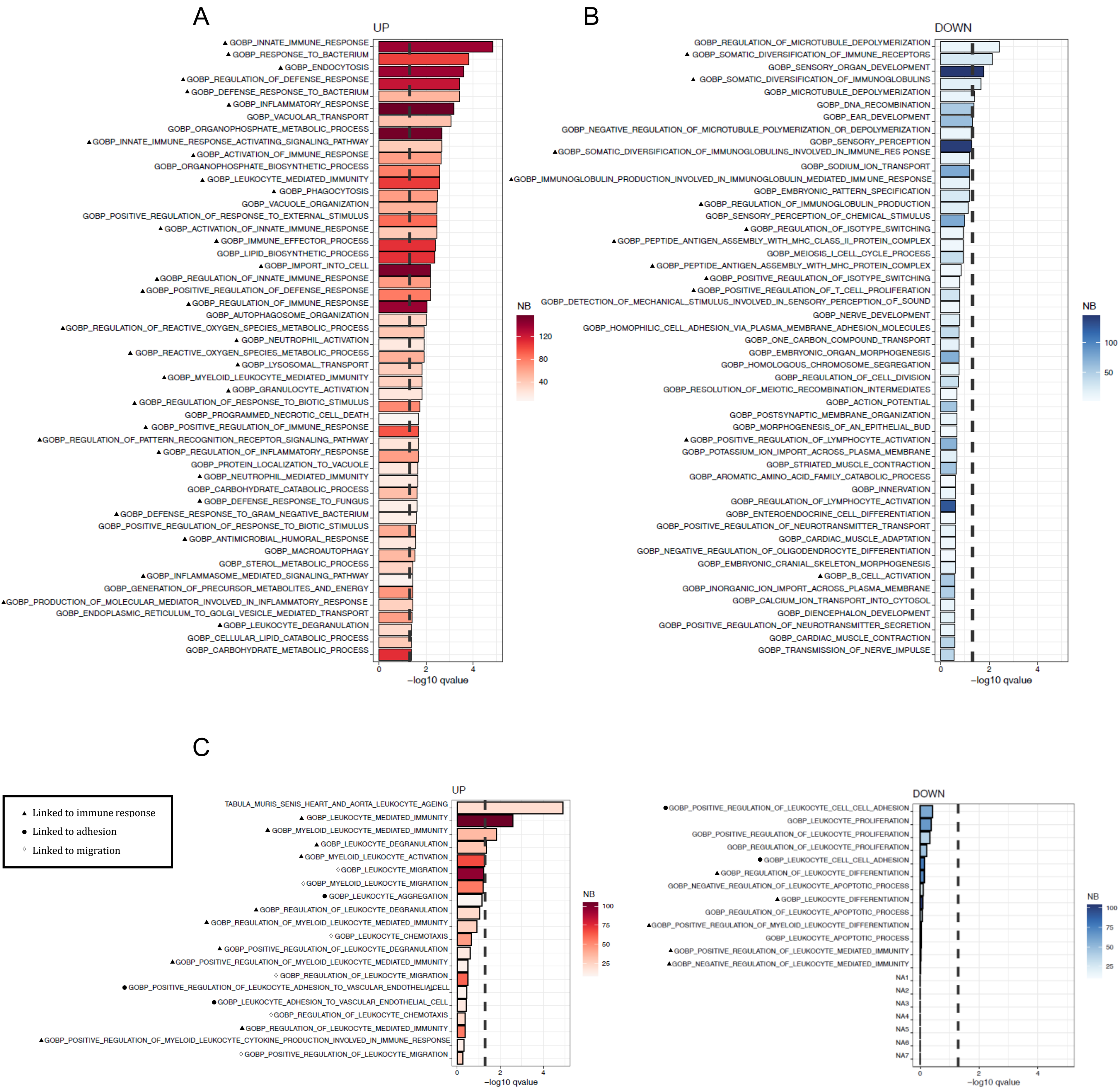


**Supplementary Figure 3 | Tumor-derived factors reprogram bone marrow-derived cells (BMDCs) toward immune-associated pathways.** FC1245 pancreatic cancer cells were orthotopically injected into mice. At 14 dpi, BMDCs were isolated from tibia and femur and RNA was subjected to bulk RNAseq (N=5 per group). **A,B** Bar plots showing the top 50 **A** upregulated and **B** downregulated Gene Ontology (GO) biological processes in BMDCs from tumor-bearing versus control mice. GO terms are ranked by $-\log_{10}$(adjusted P-value); color scale indicates the number of genes in the leading edge of GSEA. **C, D** Top 20 **C** upregulated and **D** downregulated leukocyte-related GO terms in tumor-bearing versus control BMDCs.

A

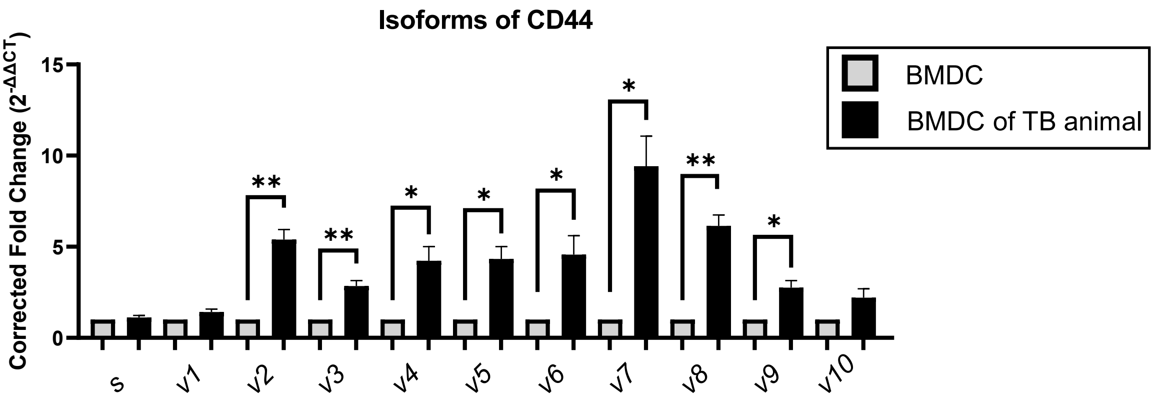


**Supplementary Figure 4 | Expression of variable isoforms of CD44 in bone marrow-derived cells (BMDCs) in response to the primary tumor.** FC1245 cells were injected into the pancreas of mice. At 14 dpi, BMDCs were isolated from tibia and femur. Relative expression of CD44 isoforms in BMDCs, normalized to Cd44pan (total CD44) and compared to controls. N = 5.

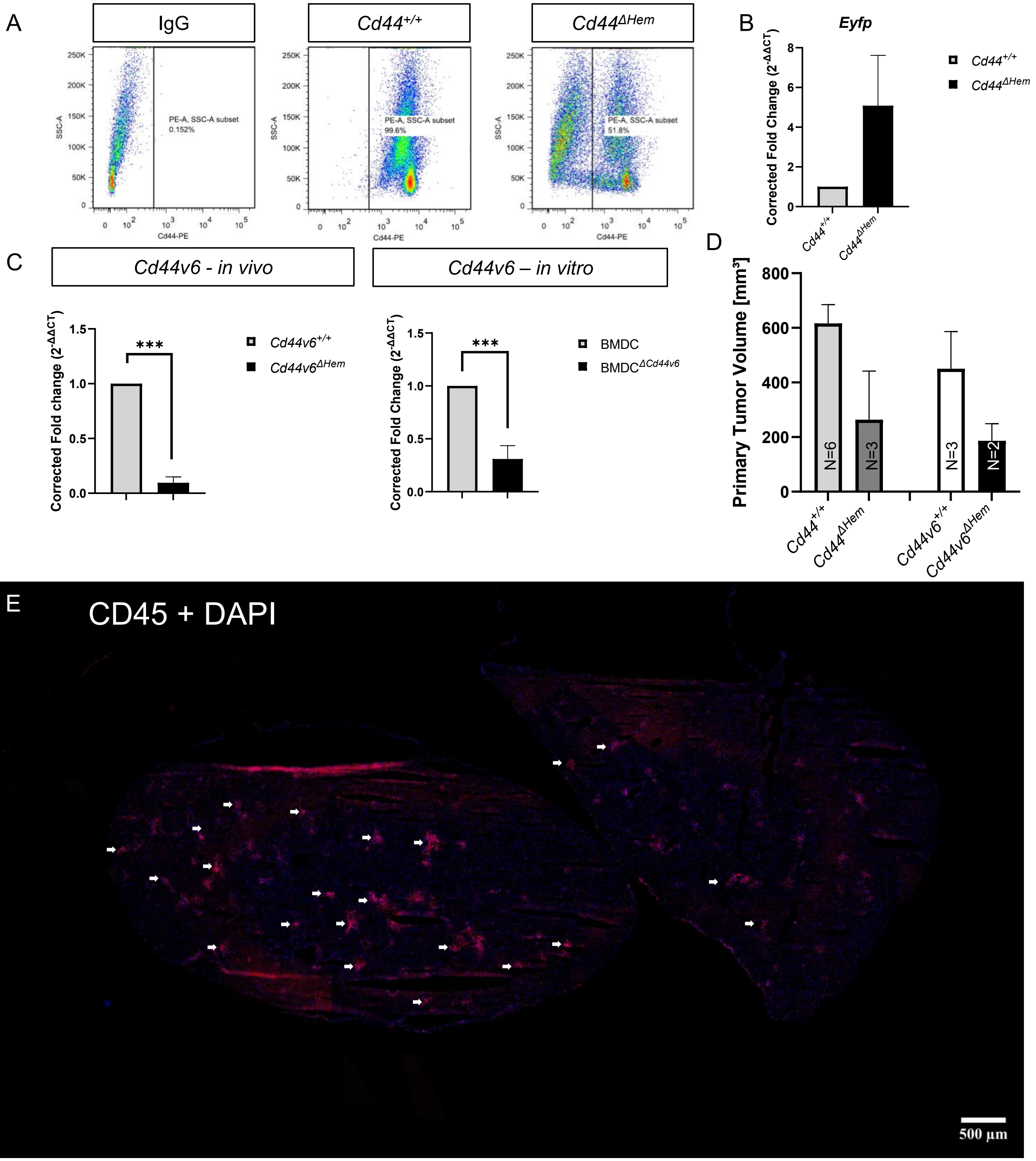


**Supplementary Figure 5 | Hematopoietic-specific deletion of Cd44 and Cd44v6 using a tamoxifen-inducible Cre/LoxP system.** A CD44 surface expression on BMDCs from *Cd44*$^{+/+}$ and *Cd44*$^{\Delta Hem}$ animals after 7-day conditioned medium (CM) of a syngeneic co-culture + 4-hydroxytamoxifen (4OHT) treatment, assessed by flow cytometry using anti-CD44-PE (clone IM7) or isotype control. B Expression of EYFP reporter calculated as fold change (2–ΔΔCT) normalized to *Gapdh*. C *In vivo* knockout validation: *Cd44v6*$^{+/+}$ and *Cd44v6*$^{\Delta Hem}$ mice received tamoxifen for 5 days, followed by orthotopic injection of FC1245 cells on day 8. At 14 or 21 dpi, liver, tumor, and BMDCs were collected. *Cd44v6* expression in BMDCs was analyzed by RT–qPCR. *Cd44v6* mRNA expression in BMDCs from *Cd44v6*$^{+/+}$ and *Cd44v6*$^{\Delta Hem}$ mice treated in vitro with CM and 4OHT; fold change (2–ΔΔCT) normalized to *Gapdh* and control BMDCs. In vivo: N = 3, In vitro: N = 5. D Measurement of primary tumor volume at 14 dpi - 21 dpi. E Exemplary overview image of CD45+ cell clusters in the liver of tumor bearing mice. Confocal image taken with Leica microhub system (MICA).

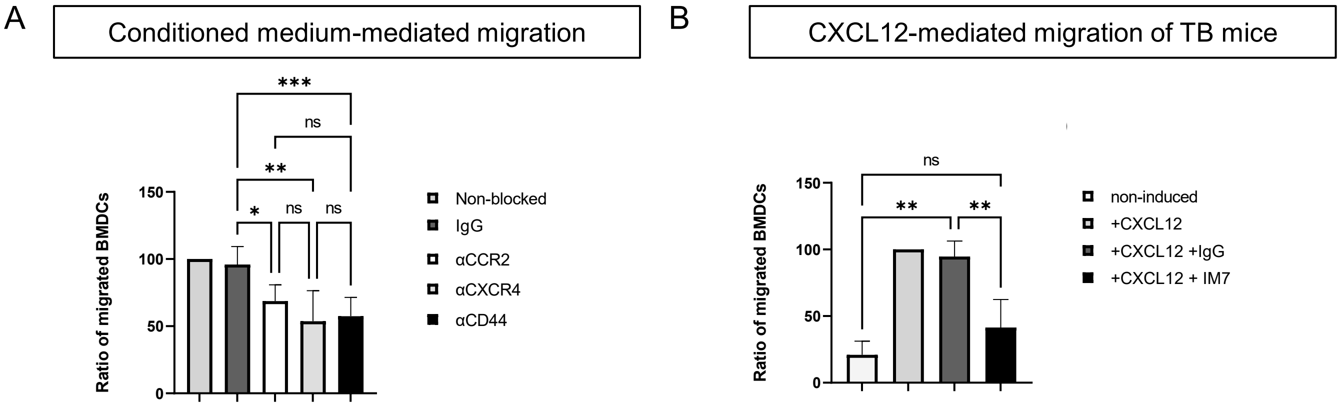


**Supplementary Figure 7 | Bone marrow-derived cells (BMDCs) were isolated from tibia and femur of healthy C57BL/6 mice. A** FC1245 PDAC cells were co-cultured with syngeneic immortalized pancreatic stellate cells (imPSC) and bone marrow-derived macrophages (BMDM). The resulting conditioned medium (CM) was applied to BMDCs. BMDCs were cultured for 3 days and subsequently seeded in transwell inserts. BMDCs were allowed to migrate for 5 hours in the presence of tumor-derived CM. Migrated cells were counted. BMDCs were treated with inhibitors against CCR1/2/5 (BX471), CXCR4 (AMD3100), CD44 blocking antibody (clone: IM7), or IgG control. **B** BMDCs were isolated from tumor-bearing animals at 14 dpi. BMDCs were seeded in a transwell, with the administration of CXCL12 in the lower well as indicated. After 5 hours, migrated cells in the lower well were counted. Statistical significance was determined using the ordinary one-way analysis of variances (ANOVA) and Holm-Šídák's multiple comparisons post-hoc test; ns=not significant, *p-value<0.0332; **p-value<0.0021; ***p-value<0.00021; ****p-value<0,0001.

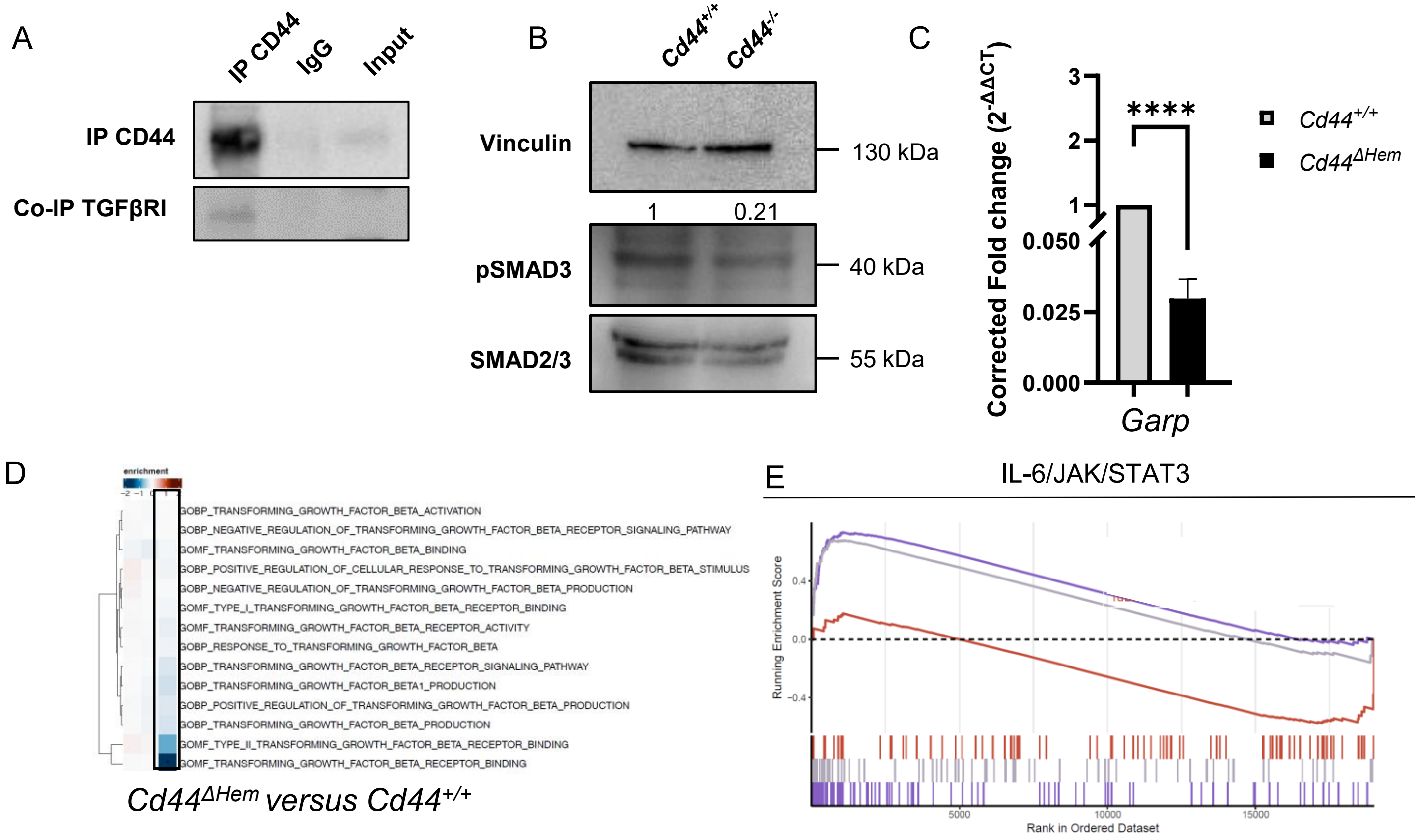


| | p-value | Adjusted p-value |
|---|---|---|
| Tumor-bearing *Cd44*$^{+/+}$ vs non-tumor-bearing | 5.6e-05 | 7e-04 |
| Tumor-bearing *Cd44*$^{\Delta Hem}$ vs non-tumor-bearing | 0.0017 | 0.021 |
| Tumor-bearing *Cd44*$^{\Delta Hem}$ vs tumor-bearing *Cd44*$^{+/+}$ | 0.043 | 0.18 |

**Supplementary Figure 8 | CD44 forms a complex with TGF-β receptor I (TGFβRI) and participates in ligand binding on BMDCs. A** Co-immunoprecipitation (Co-IP) of CD44 and TGFβRI in BMDCs. BMDCs isolated from C57BL/6 animals were cultured with conditioned medium from a co-culture of FC1245 PDAC cells, imPSCs, and BMDMs. Protein lysates were incubated with a CD44 antibody (clone: KM201) or IgG control, and the complexes were precipitated using Protein G agarose beads. Immunoprecipitation was followed by western blotting, detecting CD44 and TGFβRI. Input represents non-precipitated lysate. **B** BMDCs were isolated from *Cd44*$^{\Delta Hem}$ and *Cd44*$^{+/+}$ mice and cultured for 7 days in conditioned medium of the syngeneic co-culture. BMDCs were treated with 4OHT daily. Whole protein lysates of BMDCs were subjected to western blot analysis for pSMAD3, with SMAD3 and vinculin as a loading control. **C** BMDCs from *Cd44*$^{\Delta Hem}$ and control (*Cd44*$^{+/+}$) mice were isolated at 14 dpi after intraperitoneal tamoxifen treatment and orthotopic FC1245 injection. mRNA levels were analyzed by real-time qPCR, with *Garp* expression normalized to *Cd44*$^{+/+}$. **D** RNAseq heatmap of enriched terms related to TGF-β signaling in BMDCs from *Cd44*$^{\Delta Hem}$ versus *Cd44*$^{+/+}$ animals. Color code represents the strength of enrichment (−log10 adjusted p-value for upregulated, +log10 adjusted p-value for downregulated terms). **E** Enrichment plot of the IL-6/JAK/STAT pathway. Enrichment score (y-axis) reflects the running sum statistic from left (most upregulated) to right (most downregulated). The peak of each curve represents the point at which maximum enrichment score (ES) was reached. Each curve corresponds to a distinct comparison (color coded).
